# Modeling development of tertiary lymphoid structures in pulmonary tuberculosis by 3D profiling and trajectory analysis

**DOI:** 10.64898/2026.06.02.725201

**Authors:** Emmanuel C. Ogbonna, Soheil R. Talemi, Zoltan Maliga, Ziyuan Zhao, Clarence Yapp, Shishir M. Pant, Yi Daniel Lu, Jeremy Muhlich, Andréanne Gagné, Amanda J. Martinot, Shannon Coy, Sandro Santagata, Angela R. Shih, Isaac H. Solomon, Threnesan Naidoo, Bree B. Aldridge, Adrie J.C. Steyn, Peter K. Sorger

## Abstract

Tertiary lymphoid structures (TLSs) are sites of immune organization in peripheral tissues that arise from chronic inflammation. They play important roles in infection control and cancer but the mechanisms controlling their formation remain only partly understood. Here, we combine high-plex imaging, serial-section 3D reconstruction, and optimal transport trajectory modeling to reconstruct TLSs in human lungs infected with *Mycobacterium tuberculosis*. We find that the extended and irregular shape of TLSs is poorly captured by 2D histopathology or spatial profiling, making assessment of developmental stage (early, primary, or secondary) error prone. In contrast, modeling TLS development in 3D using optimal transport reveals two trajectories that differ in the timing and coordination of follicular dendritic cell association, germinal center consolidation, and position within the tissue. These findings highlight the value of volumetric analysis in understanding immune organization and provide new insights into TLS biology.

**Graphical Abstract:** 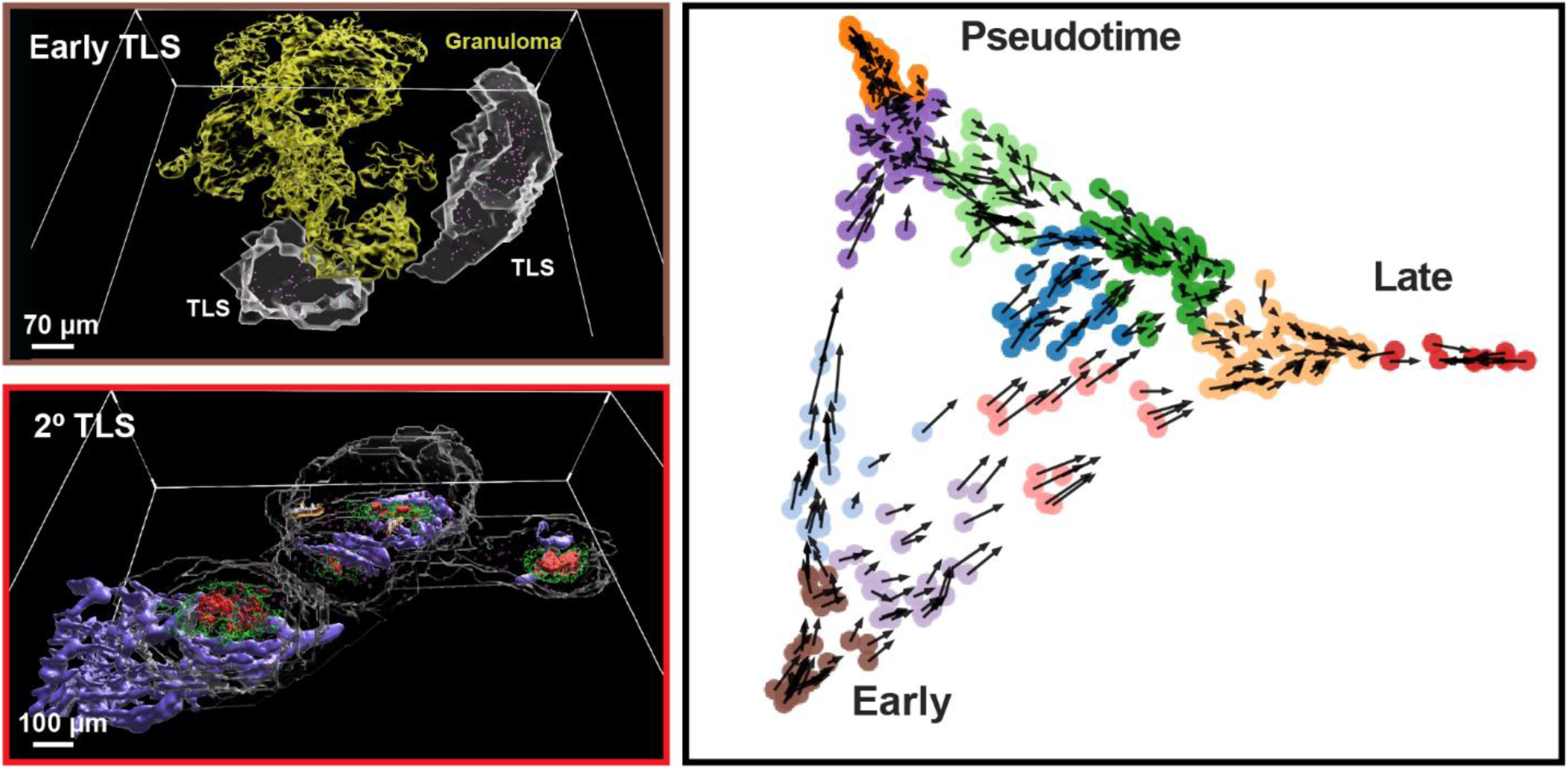

**Highlights:**

- 3D reconstruction of TB-infected lung reveals TLSs to be extended, heterogeneous structures that are frequently misclassified in 2D
- TLS maturation follows continuous trajectories rather than discrete histological stages
- Optimal transport modeling identifies distinct maturation paths with divergent follicular dynamics
- Germinal center organization and Tfh-FDC interactions emerge along maturation trajectories
- TLS maturation states are shaped by spatially localized microenvironmental niches.

## INTRODUCTION

Tuberculosis (TB) is the leading cause of mortality from a single infectious disease worldwide, resulting in ∼10.8 million infections and 1.25 million deaths in 2023.^1,2^ Reducing the burden of disease is challenged by drug resistance, variable vaccine efficacy, and incomplete understanding of TB immunopathology.^3^ Infection with *Mycobacterium tuberculosis* (*Mtb*), has variable immune and pathological effects across patients and tissues.^4–6^ Only 5-10% of individuals exposed to *Mtb* develop active disease with others instead developing sub-clinical TB, a portion of which represents latent infection.^2,7–9^ Latent TB can reactivate into an infectious state, particularly in immunocompromised individuals.^10,11^ The bacterial and host factors that govern different responses to infection and the efficacy of treatment or vaccine prophylaxis are not fully understood.

Granulomas are characterized by collections of epithelioid histiocytes (activated macrophages) and other immune cells that aggregate in response to chronic inflammation. This results in formation of dense immune aggregates that function to isolate *Mtb* infection. The granuloma microenvironment (GME)^12–14^ is a hallmark of TB and plays a key role in bacterial clearance, immune containment, establishment of latency, and responsiveness to treatment.^15–18^ Depending on the stage of infection, granulomas can be primarily cellular or necrotic. Ectopic granuloma-associated lymphoid tissues (GrALTs) form in association with granulomas and represent TB-specific instances of tertiary lymphoid structures (TLSs).^12,19–24^ TLSs are sites of B and T cell programming and amplification outside of immune tissues and are linked to enhanced immune protection, particularly following vaccination with attenuated *Mtb* strains or primed immune cells.^21,25–27^ For example, macaques vaccinated with the experimental *MtbΔsigH* vaccine exhibit increased TLS formation and T cell recruitment, and improved infection control relative to animals vaccinated with the traditional Bacille Calmette-Guérin (BCG) vaccine,^21^ a finding mirrored in murine models.^25,26^ Thus, understanding the biology of TLSs and peripheral immunity in TB is likely to play an important role in advancing new TB vaccines and host-directed therapies.^28^

Unlike primary and secondary lymphoid organs, which are developmentally programmed for antigen presentation and clonal selection,^29,30^ TLSs are non-encapsulated immune complexes that form *de novo* due to chronic inflammation^31–34^ arising from infection, cancer, and tissue repair.^21,23,24,34–37^ Specifically in the lung, the lymphoid cells normally found in association with bronchi and bronchioles expand dramatically in response to *Mtb* infection and give rise to bronchus-associated lymphoid tissue (BALTs)^38^ that are the counterpart of TLSs found in other peripheral tissues. Most (perhaps all) TLSs in *Mtb*-infected lungs are BALTs and a subset of these are also GrALTs^12^ (for simplicity we will refer to these related structures in aggregate simply as TLSs). Architecturally, TLSs are comprised of B and T cell zones, specialized vasculature (e.g. high endothelial venules; HEVs), and networks of follicular dendritic cells (FDCs), that organize the germinal centers (GC) involved in antigen presentation and B cell activation.^23,24,31,33^

Individual TLSs vary with respect to these biological and structural features depending on their state of maturation: early TLSs are loose lymphocytic aggregates lacking expression of CD21 (complement receptor 2) and CD23 (an IgE receptor) on FDCs and also some mature B cells; primary TLSs contain CD21⁺ FDC networks (and are thus CD21⁺ CD23⁻) and secondary follicle-like TLSs have CD21⁺ CD23⁺ FDC networks with GC morphology and activity. This has given rise to a widely used early-primary-secondary (EPS) classification scheme that has been further refined based on 2D spatial profiling as well as dissociative bulk and scRNA sequencing.^39–44^

A key question in TB immunopathology is precisely how TLSs develop and mature and what roles they play in *Mtb* clearance, containment, and persistence. Similar questions arise in diseases such as cancer, in which TLSs are key features of the tumor microenvironment. We had previously established that TLSs in colorectal cancer are complex 3D structures several hundred microns in extent,^45^ but due to limitations in markers and imaging methods neither we, nor other groups, have as yet investigated the implications of this 3D organization for understanding TLS development; in this manuscript, we address this gap in knowledge. We use 3D serial section reconstruction of high-plex CyCIF images of lung tissue from individuals infected with *Mtb* to identify a set of ∼900 B cell-rich immune complexes with the properties of TLSs and ranging in size from ∼10^2^ to over 10⁵ cells. When only 2D sections from these TLSs were scored, misclassification rates (in the EPS framework) exceeded 50% relative to the 3D ground truth. Even with 3D data, EPS staging poorly represented the state of TLS maturation with respect to key immunological and molecular characteristics. In contrast, a trajectory-based approach based on optimal transport (OT)^46^ was effective at modeling TLS maturation in 3D as a continuous developmental process, providing insight into TLS biology not readily discernable in 2D.

## RESULTS

### 3D reconstruction and multimodal profiling of TLSs in TB-infected lung

Spatial profiling of untreated *Mtb*-infected lung tissue was performed by 30 to 55-plex whole-slide cyclic immunofluorescence (CyCIF)^47^ or hematoxylin and eosin (H&E) staining of 5 µm formalin-fixed paraffin-embedded (FFPE) sections. CyCIF markers were chosen to distinguish immune, epithelial, and endothelial cell types, specify cell states such as proliferation and activation, and confirm the presence of *Mtb* (based on Ag85b staining). These data were supplemented by thick section 3D CyCIF using confocal microscopy^48^ and GeoMx spatial transcriptomics on selected tissue sections. A board-certified thoracic pathologist reviewed and annotated H&E images to guide interpretation of CyCIF data.

In a pilot study of 10 serial sections from a representative tissue block, inspection of H&E and CyCIF images revealed the presence of granulomas flanked by lymphocyte-rich aggregates positive for the B cell marker CD20 and T cell marker CD3 (**Figures 1A-B**). Spatial transcriptomic profiling (GeoMx) of CD20-positive features confirmed the expression of chemokines associated with TLS maturation, including CXCL13, CCL19, CCL21, CCL5, and CXCL9.^49^ (**Figures S1A-C; Table S1**). For these and other genes, trajectory analysis yielded pseudotime axis that recapitulated known features of TLS maturation including increasing expression of genes specific to B cells, the CXCL13 cytokine and its receptor CXCR5, and markers of FDC differentiation (**Figures S1E-G**). We conclude that TLSs in our dataset closely resembled immune structures previously observed in cancer, TB,^12^ and a range of peripheral tissues subject to chronic infection.^37^

**Figure 1.**
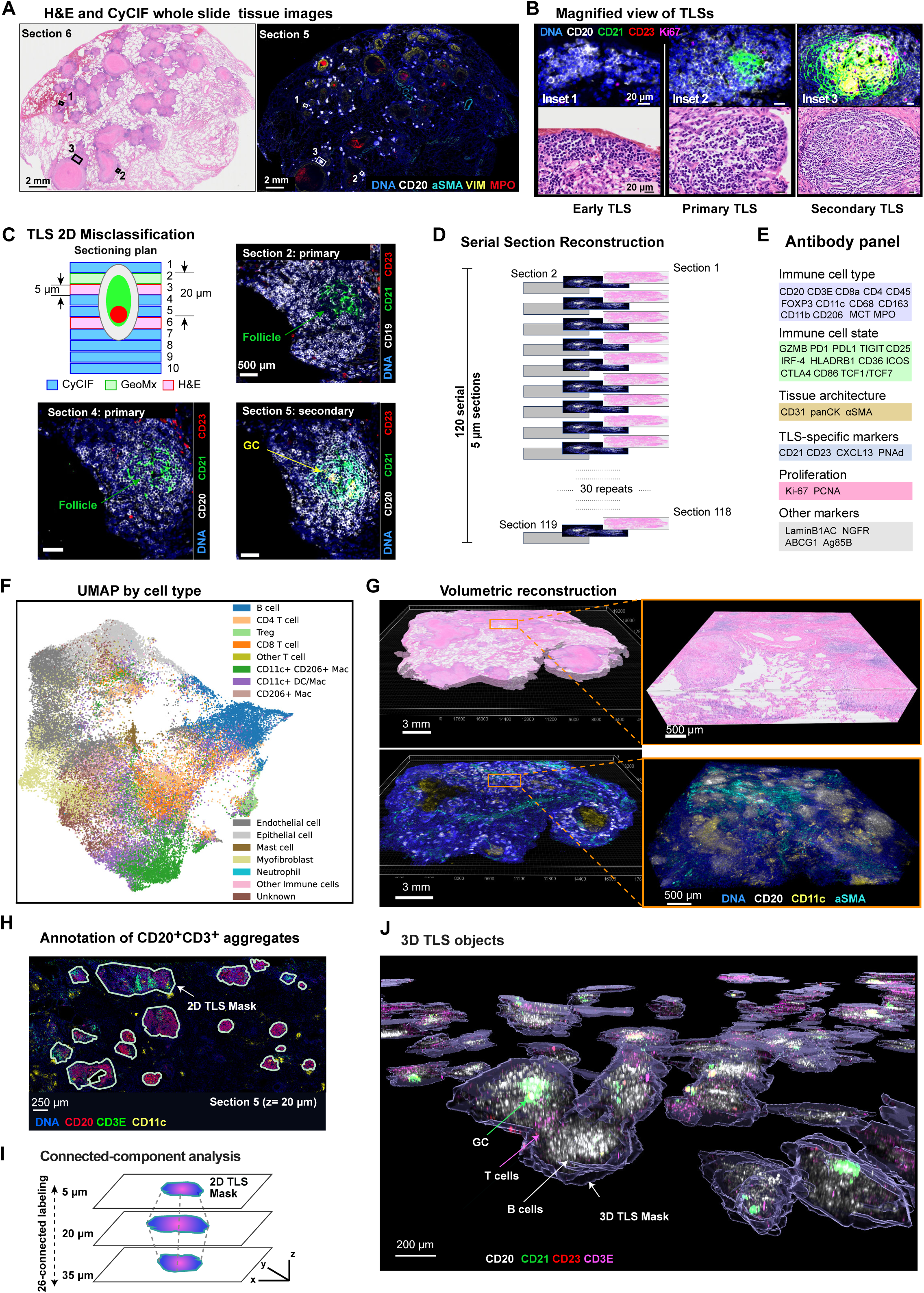
Multimodal spatial profiling and three-dimensional reconstruction of tertiary lymphoid structures in pulmonary TB. **(A)** Representative whole-slide 2D CyCIF and H&E images from TB-infected human lung tissue (Patient1_B5), highlighting granulomas and surrounding lymphoid aggregates (DNA stain with Hoechst 33342, blue; CD20, white; αSMA, cyan; vimentin, yellow; myeloperoxidase, red). **(B)** Representative TLSs illustrating canonical histological classes defined in 2D, with distinct spatial organization of CD20 (white), CD21 (green), CD23 (red), and Ki-67 (magenta). **(C)** Sectioning-dependent TLS misclassification. Top left: Schematic showing Patient1_B5 tissue sections used in pilot study. Sections are colored by imaging modality. CyCIF was performed on sections 2, 4-5, and 7-10, ORION immunofluorescence on section 1 (both modalities colored light blue),^112^ and H&E on sections 3 and 6 (light pink). On section 2, CyCIF was done following GeoMx spatial transcriptomics (light green). A single TLS exhibits CD21⁺CD23⁻ primary features in sections 2 (top right) and 4 (bottom left) but displays CD21⁺CD23⁺ secondary organization in the adjacent section 5 (bottom right). **(D)** Experimental design for 3D reconstruction of TLSs from the Patient1_B4 tissue block. A total of 120 consecutive 5-µm FFPE sections were collected, spanning ∼600 µm; 40 sections were used for H&E imaging and adjacent sections were selected for 39-plex whole-slide CyCIF imaging, with interleaved section skipping. **(E)** CyCIF antibody panel used for 3D reconstruction experiment, capturing markers relevant to TLS architecture and function. The complete antibodies list is provided in **Table S3**. **(F)** Uniform manifold approximation and projection (UMAP) plot of single-cell data from CyCIF imaging (Patient1_B4_Section 2), colored by cell phenotype. UMAP was generated from randomly sampled 50,000 single cells. See also **Figure S2D**. **(G)** Registered H&E (top left) and CyCIF (bottom left) image volumes. Inset from bottom: region revealing multiple TLSs (CD20, white) associated with granulomas (CD11c, yellow), and myofibroblasts in proximal blood vessels (cyan); Inset from top: corresponding H&E view. **(H)** Annotation of CD20⁺CD3⁺ TLS regions in 2D CyCIF images across the reconstructed stack, yielding ∼6,000 TLS regions of interest (ROIs) (DNA, blue; CD20, red; CD3E, green; CD11c, yellow). See also **Figure S3B**. **(I)** 3D reconstruction of TLS ROIs by 26-connected-component labeling across serial sections. Reconstructed TLS volumes were curated to generate a final set of discrete 3D TLS objects for downstream spatial and quantitative analyses. **(J)** Representative 3D TLS objects.

However, when a single TLS was followed through adjacent tissue sections, the assignment of early, primary or secondary maturation state varied with the section analyzed. For example, the TLS in **Figure 1C** was CD21⁺CD23⁻ in Sections 2 and 4 (corresponding to a primary TLS) but CD21⁺CD23⁺ in Section 5 (consistent with a secondary TLS). Separately, serial-section CyCIF of non-small cell lung cancer, another pulmonary disease, revealed similar limitations in 2D assignment of TLS state (**Figure S2A**), as well as a broadly similar distribution of TLS sizes and morphologies.^50^ Thus, the problem of TLS mis-annotation is not restricted to TB lung. This makes intuitive sense, since TLSs are complex 3D structures that are conventionally viewed in 2D cross-section^45^ but the impact of mis-annotation has not generally been considered in the analysis of TLS development and function.

We therefore performed a volumetric reconstruction of TLSs in a 2 cm x 1.5 cm x 600 µm FFPE block of TB-infected lung tissue using a “skip-2” scheme involving 120 consecutive 5 µm sections with every third slide undergoing 39-plex CyCIF whole-slide imaging (**Figures 1D-E; Table S2**). Adjacent slides were H&E stained and intervening slides held in reserve (**Figure 1D**). Across 40 CyCIF WSIs, a total of ∼47 million cells were assigned to 13 immune and stromal cell types (**Figures 1F**) and a range of functional cell states (**Figures S2C-D**). Reconstruction of a 3D image from tissue sections required several technical innovations: the CODA method^51^ used to generate 3D data of human pancreas provided inspiration but was not effective with lung tissue, probably because lung is less rigid with many acellular spaces (e.g. alveoli); these features complicate the alignment procedure. We therefore developed a multistage alignment pipeline based on Advanced Normalization Tools (ANTs),^52^ a software environment widely used for radiological (MRI and CT) imaging, particularly in neuroscience.^53^ Our implementation used nuclei to guide rigid, affine, and nonlinear SyN transformations and reconstruct skip-2 CyCIF images at full resolution (**Figure S3A**; see Methods).^54,55^ H&E sections were then co-registered using elastix (**Figure 1G**).^56^

### 2D imaging substantially underestimates the proportion of secondary TLSs

Regions of tissue rich in B and T cells (CD20⁺or CD3E⁺) were identified using a standard pixel classifier in QuPath^57^ to generate ∼6,000 2D TLS ROIs (∼110 to 190 per section; **Figure S3B, S1D**); volumetric reconstruction of these ROIs using connected-component analysis consolidated them into a total of 906 3D lymphocyte-rich structures. Each structure was defined by a mesoscale segmentation mask and corresponding single cell data; CyCIF and H&E images were also compared manually and computationally (**Figures 1H-J, S3C**). To identify TLSs that were interrupted by tissue cutting planes, we used a bounding box metric: TLSs were classified as “complete” if their bounding box lay entirely within the reconstructed volume, or “partial” if it intersected volume boundaries.^58^ This yielded 640 complete TLS objects (71% of the total; **Figure S3D**). Quality control of these 640 structures using Cylinter^59^ to identify artifacts of tissue processing and staining resulted in a final set of 470 TLSs suitable for detailed analysis.

Expression of CD21 and CD23 within each TLS volume was used to stage each structure as early, primary or secondary in accordance with previously described 2D imaging approaches.^44^ We found the 470 3D structures were mostly (69%) secondary with fewer primary (16%) or early TLSs (15%; **Figure S4A**). This is different from the 36% secondary, 41% primary and 23% early distribution estimated from 2D images (**Figures S4B**). Ultrastructural variants, such as multifocal primary and secondary TLSs have 2 or more follicles or GCs were also common (93/470; 20%) and were largely undetected in 2D images. Thus, conventional 2D imaging substantially overestimated the proportion of early and primary TLSs, underestimated the proportion of secondary TLSs, and missed multifocal TLSs.

### Organization of follicular dendritic cells and germinal centers

Most of what is known about the 3D architecture of lymphoid structures derives from serial-section (“whole-volume”) H&E imaging of lymph nodes supplemented by low-plex light sheet fluorescence^60^ or confocal imaging^61^ of murine lymph nodes. Much less is known about the overall architecture of human TLSs, and the data that are available derive from tumors,^37^ not infected tissues. In our dataset, TLSs varied by three orders of magnitude in cell number and size, from ∼10^2^ to >10⁵ cells (**Figures S3E, S4C-D; Video S1**), 10 to 30-fold more than found in a single 2D section. 3D morphologies and molecular characteristics varied widely (**Figures S4E-F**). For example, early TLSs 594 and 597 were compact ellipsoidal GrALTs adjacent to lymphocytic cuffs (loosely organized concentrations of T cells surrounding granulomas; **Figure 2A**). Primary TLS 321 wrapped around a granuloma (**Figure 2B**), multi-focal secondary TLS 80 had four well-formed CD21⁺ CD23⁺ GCs (**Figure 2C**); and TLS 154 had two GCs, with a prominent intervening panCK⁺ bronchiole (**Figure 2D**).

**Figure 2.**
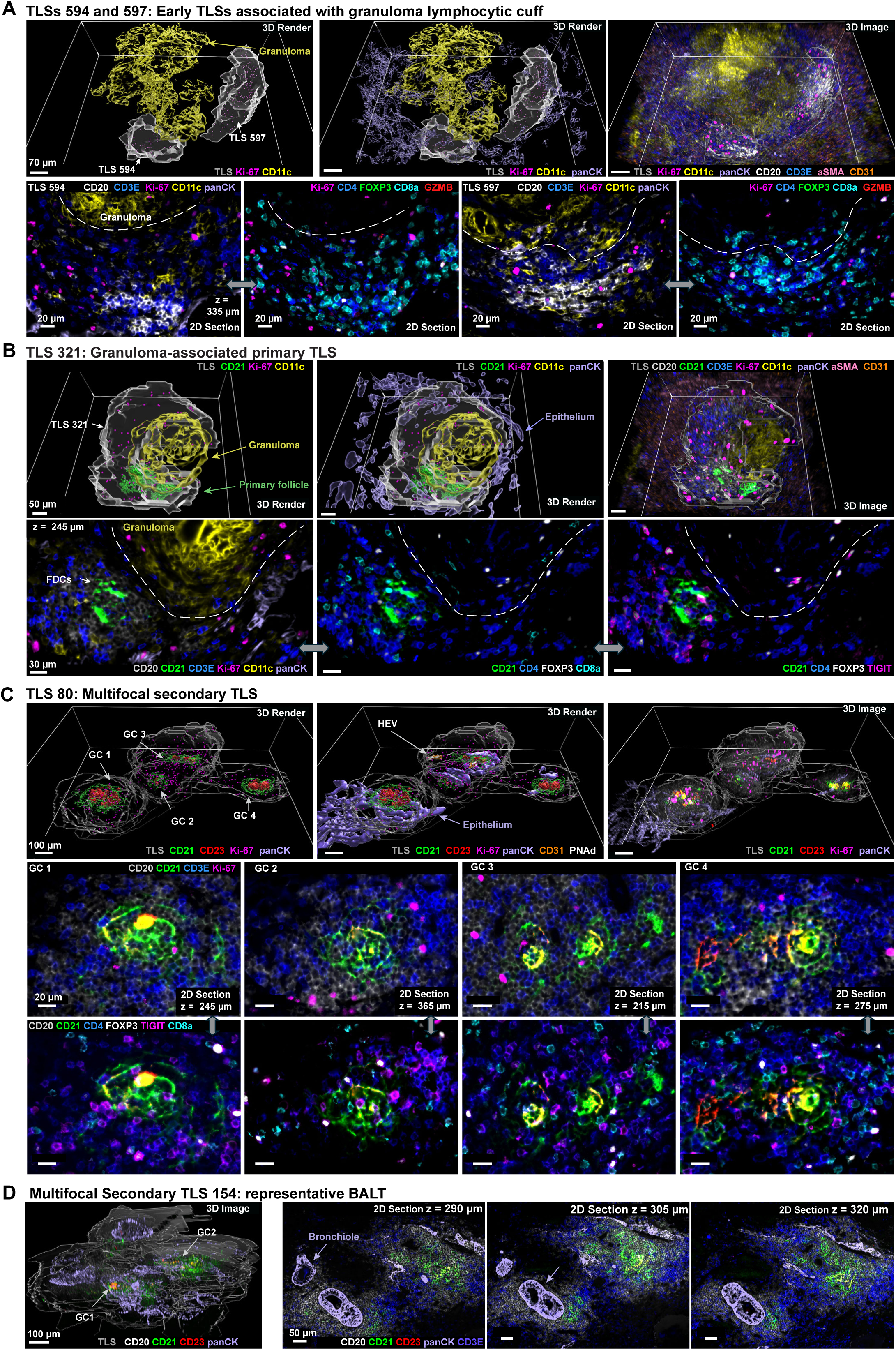
Three-dimensional microarchitecture of TLS maturation stages in pulmonary TB. **(A)** Representative early TLSs reconstructed in 3D from Patient1_B4, located at the granuloma periphery. These structures comprise mixed CD20⁺ B cells and CD4⁺/CD8⁺ T cells but lack follicular dendritic cell (FDC) development (CD21⁻CD23⁻). **(B)** Primary TLS exhibiting compact lymphoid organization with emergence of a CD21⁺ FDC network (but absence of CD23 expression) and dispersed Ki-67⁺ proliferating lymphocytes which indicate early follicular structuring. **(C)** Secondary TLS displaying multifocal germinal center-like organization characterized by CD21⁺CD23⁺ FDC networks, Ki-67⁺ proliferative activity, and TIGIT⁺ T follicular helper (Tfh) cells. Also highlighted were PNAd⁺CD31⁺ HEVs. See also **Figures S4, S5,** and **5**. **(D)** Representative bronchus-associated lymphoid tissue (BALT).

Across all TLSs, B cells were the majority cell type, followed by CD11c⁺ myeloid cells, CD4⁺ T cells and vascular endothelial cells (**Figures 3A, S4E**). PanCK-positive cells were associated with nearly all TLSs (∼95%) either as well-formed bronchioles or inflamed and disorganized epithelia (**Figures 2C-D**). Linear regression of cell-type abundances with TLS size revealed enrichment for B cells, T cells and CD11c⁺ DC/macrophages, and depletion of neutrophils, mast cells, and CD206⁺ macrophages (**Figure 3B**). As TLSs matured and grew, they also became increasingly zonated (shown schematically in **Figure 3C** and with examples in **Figure 3D**) with segregation of B from T cells (blue and orange, **Figure 3D**), assembly of a follicular FDC network (green) surrounded by B cells, and finally GC formation (red) within the follicle. To quantify zonation, we applied a microregional labeling strategy that accounted for irregular compartment shapes, uneven distribution of FDCs, and the challenge of accurately segmenting highly elongated FDCs from each other. TLS outer boundaries were identified using a pixel classifier based on patterns of CD20 and CD3 staining (as described above). Within these boundaries, spatially contiguous “seed” domains were identified using *k-*nearest neighbor (kNN) graphs (**Figure 3E**). Seeds were expanded radially to incorporate neighboring cells and generate continuous domain labels, irrespective of intermixing of other types of cells. Labels were applied hierarchically: CD21⁺ CD23⁺ FDC zones were defined as GCs. CD21⁺ FDC zones as follicles (primary when a GC was absent and secondary when it was present), and CD20⁺ enriched regions as B cell zones. By definition, B cell zones excluded the follicle and GC, and the follicle excluded the GC (**Figure 3C**). Zonal location and markers were also combined to identify key cell types such as T follicular helper and regulatory cells (Tfh and Tfr; **Figures S5A-D**). This provided a robust and scalable approach to analyzing TLS zones and their cellular constituents in 3D.

**Figure 3.**
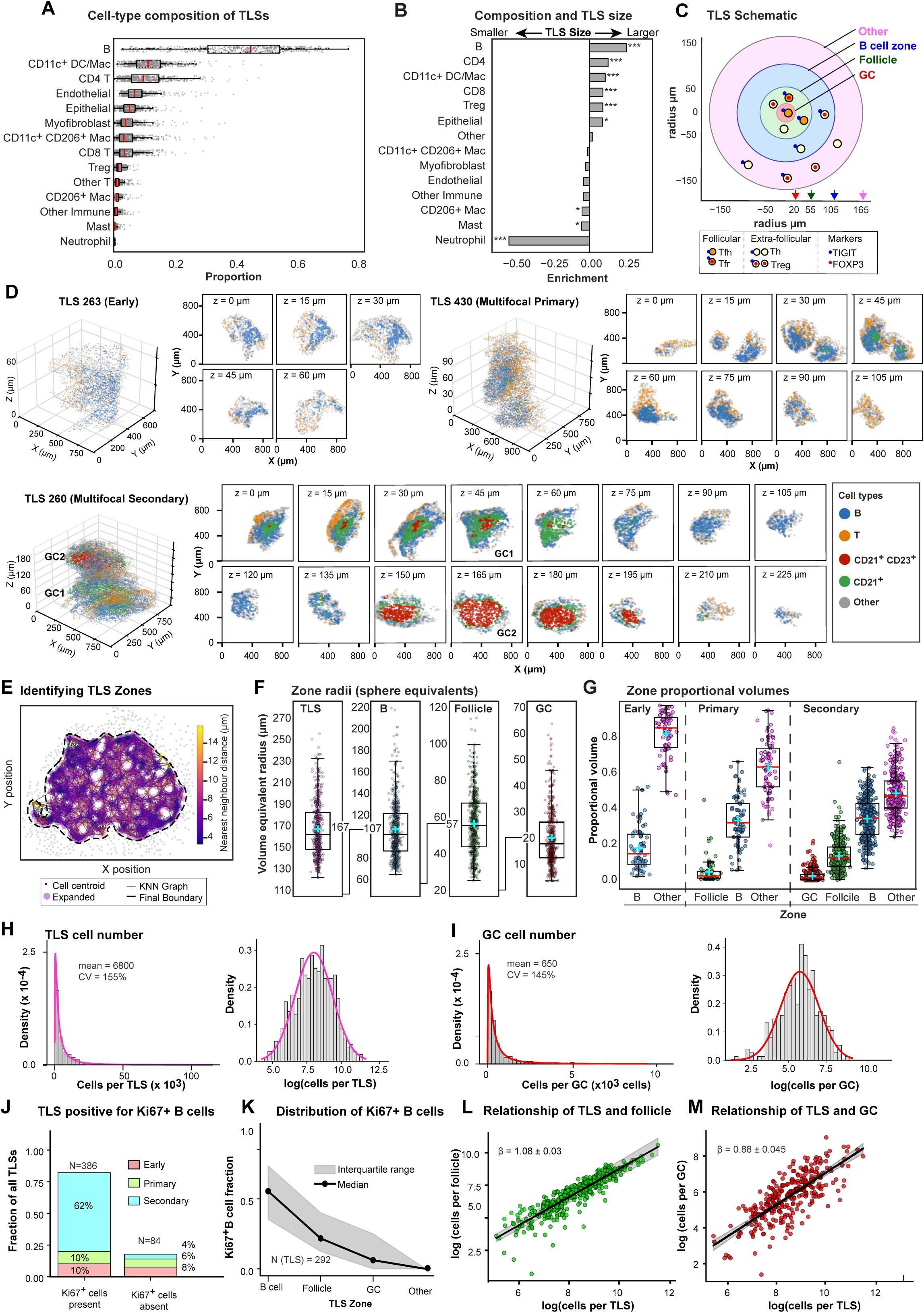
Cellular composition, follicular organization, and scaling properties of TLSs. **(A)** Compositional analysis of reconstructed 3D TLSs. Distribution of cell-type fractions per TLS across all reconstructed 3D TLSs, with the 5th–95th percentile range (black horizontal line), interquartile range (light red line inside box), and mean (red vertical line) indicated for each cell type. **(B)** Linear regression analysis of TLS composition with TLS size as a covariate. Cell types are ranked by β coefficients from linear models relating CLR-transformed cell-type abundance to log-transformed total cell count per TLS. CLR values were calculated from TLS-level cell-type counts with a pseudocount of 1. β coefficients quantify the association between relative cell-type abundance and TLS size. Positive values indicate relative enrichment in larger TLSs, whereas negative values indicate relative depletion. Asterisks indicate nominal regression significance based on two-sided p values (*p < 0.05, **p < 0.01, *p < 0.001). **(C)** TLS schematic showing concentric shells based on mean volume-equivalent radii of TLS zones. The spatial positioning of follicular and extrafollicular T cells are also indicated. **(D)** 3D and 2D renderings of representative TLSs illustrating spatial segregation of CD20⁺, CD21⁺, and CD23⁺CD21⁺ zones in early, multifocal primary, and multifocal secondary TLSs. **(E)** Representative 2D k-nearest neighbor (kNN) graph used for zone construction within a TLS section. Nodes represent individual cell centroids, edges denote local kNN relationships, and the dashed contour indicates the expanded boundary generated from the connected graph structure. Example shown corresponds to a CD21⁺CD23⁺ domain from TLS260 at z = 260 µm (Section 53). **(F)** Distribution of volume-equivalent radii of germinal center (GC), CD21⁺ follicle, CD20⁺ B cell zone, and “Other” zone (rest of TLS). Volume-equivalent radii were calculated assuming spherical geometry. For visualization, compartment radii are displayed on cumulative y-axes generated by adding the mean radii of the preceding compartment(s), reflecting the nested TLS architecture. Elbow connectors indicate the resulting cumulative mean radii: 20.21 µm (GC), 56.52 µm (GC + follicle), and 106.69 µm (GC + follicle + B cell zone). The cumulative mean radius of the entire TLS was 167.19 µm. **(G)** Proportional volumes occupied by CD20⁺ B cell zones, CD21⁺ follicular dendritic cell (FDC) zones, CD23⁺CD21⁺ germinal center-like zones, and non-B cell non-follicular regions (“Other cells”) across early, primary, and secondary TLSs. Boxplots show relative zone volumes per TLS class. Cyan crosses indicate means. **(H)** TLS cell number distribution. Total cell counts per TLS, using TLSs that passed QC (N=470), plotted as a histogram on the linear scale (left) and log scale (right). The magenta curve shows the lognormal distribution fitted to the raw cell counts; the right panel shows the same fit transformed to log space. The mean cell count and coefficient of variation are shown. **(I)** Cell number distribution in germinal centers (GC). Total cell counts per GC (N=386), using TLSs that passed QC, plotted as a histogram on the linear scale (left) and log scale (right). The red curve shows the lognormal distribution fitted to the raw cell counts in GCs; the right panel shows the same fit transformed to log space. **(J)** TLS maturation stage stratified by proliferative activity (Ki-67⁺ B cell presence). Stacked bar charts showing the proportion of TLSs at each maturation stage (Early, Primary, Secondary) for TLSs with Ki-67⁺ B cells present (N=386) and absent (N=84). Early TLSs contain a CD20⁺ zone only; Primary TLSs contain a CD21⁺ follicle; Secondary TLSs contain a germinal center (GC). The percentage of each stage within each group is shown. **(K)** Distribution of Ki-67⁺ B cells across TLS zones. Median Ki-67⁺ B cell fraction (filled circles, solid line) and interquartile range (shaded area) across TLS zones (B cell, Follicle, GC, “Other”), for Ki-67⁺ B cell-positive TLSs (N=292). The fraction represents the proportion of Ki-67⁺ B cells located in each zone relative to the total Ki-67⁺ B cell count per TLS. **(L)** Relationship of TLS and follicle size. Log-log scatter plot of total cell count in follicle (CD21⁺ compartment) versus total cell count in TLS for TLSs with a detectable CD21⁺ compartment (N=386). The black line shows the power-law fit with bootstrapped 95% confidence interval (shaded, Nboots=10⁴ bootstrap samples). The allometric scaling exponent β and its bootstrap standard deviation are indicated. **(M)** Relationship of TLS and GC size. Log-log scatter plot of total cell count in GC (CD21⁺CD23⁺ compartment) versus total cell count in TLS for TLSs with a detectable GC compartment (N=296). The black line shows the power-law fit with bootstrapped 95% confidence interval (shaded, Nboots =10⁴). The allometric scaling exponent β and its bootstrap standard deviation are indicated.

A simple way to visualize relationships among TLS zones is as volumes or volume-equivalent radii (radii of spheres having the same volume as irregularly shaped TLS zones; **Figure 3F**). The volume of the B cell compartment increased monotonically from ∼18% to ∼35% from early to secondary TLSs (**Figure 3G**; blue data points) and the fraction of all other cells (primarily CD4 T cells) fell from ∼82% to ∼49% (pink data points). In primary TLSs, follicles occupied an average of 4% of the TLS volume, increasing to 14% in secondary TLSs (green data points). GCs were on average ∼5-fold smaller at 2.6% of total volume. However, zone dimensions and the ratio of follicle to GC volume varied widely, from ∼0.7 to >20-fold and this variability was observed even within a single multi-focal TLS (compare GC1 and GC2 in TLS 260; **Figure 3D**).

Despite morphological irregularity and differences in the extent of zonation, the overall size of TLSs (as measured by cell number) exhibited an excellent fit to a log-normal distribution (**Figure 3H**), a common feature of structures dependent on multiplicative processes such as cell division.^62^ In such heavy-tailed distributions, most TLSs were similar in size to the population average (∼7,000 cells) but some structures were as large as >100,000 cells (**Figure S3E**). GC size was also log-normally distributed and had a high coefficient of variation (mean = 650 cells, **Figure 3I**). These physical parameters were directly related to TLS activity: Ki-67^+^ B cells (which were primarily found in the B cell zone not follicles or GCs) were significantly more common in secondary than other TLSs (**Figures 3J-K)**. Remarkably, the numbers of cells in each TLS, follicle and GC were found to obey a power-law: *Y* = *αM*^*β*^ where *Y* is the number of cells in one zone (e.g. follicle), *M* is the number of cells in a second zone (e.g. the TLS), α is a proportionality constant, and β is the allometric exponent. Allometric scaling relationships of this type are common in physiology, most prominently in Kleiber’s law, which relates basal metabolic rate to organism size with β =0.75.^63^ In our dataset, follicle size scaled super-linearly with TLS size (β = 1.08), whereas GC size scaled sub-linearly (β = 0.88; **Figures 3L-M**). Thus, as a TLS grows, the follicle occupies an increasing proportion of the volume but GCs do not keep pace.

Consistent with the general interpretation of Kleiber’s law, this suggests that GCs become increasingly efficient at recruiting or stimulating B and T cells as a TLS grows and matures. The existence of scaling relationships among TLS components is evidence that they are linked by a common set of physiological processes linking growth and zonation.

### Optimal transport trajectory analysis reveals continuous paths of TLS maturation

The strong correlation of TLS size, composition and ultrastructural organization provided a natural framework for modeling TLS development as a continuous process using Optimal Transport (OT). OT is a form of trajectory analysis that provides a mathematical framework for mapping multiple entities – in this case, 3D clusters of immune cells – from one configuration to another at the least possible “cost.” Cost in our analysis corresponded to the number of cell birth, death, and migration (transport) events needed to transform one 3D lymphoid structure into another. OT has been previously been used to reconstruct developmental processes from single cell transcriptomic data^46,64–67^ with cost computed from gene expression dissimilarity (e.g. cosine distance of expression vectors). In contrast, OT cost in the current work has a direct relationship to the physical process of TLS assembly in 3D via division, death, and migration of individual cells. Cost was computed based on the pairwise distances between TLSs using unbalanced fused Gromov-Wasserstein optimal transport (UFGW), which jointly captures spatial and phenotypic organization (i.e. differences between immune cells of different types) while allowing for differences in cell number (**Figure 4A**). To reduce computational complexity, distances were calculated relative to a subset of anchor TLSs spanning the size range in dataset (**Figure S6A**). Distance profiles were embedded using a Deep Set encoder and visualized as a low-dimensional diffusion map, providing a compact representation of TLS relationships (**Figure S6B-C**). A flow-matching approach was then used to infer a continuous vector field over the embedding space, thereby allowing trajectories to be inferred. Pseudotime was computed for each trajectory by fitting principal curves to the corresponding TLS positions in diffusion-map space (**Figures S6C-D**).

**Figure 4.**
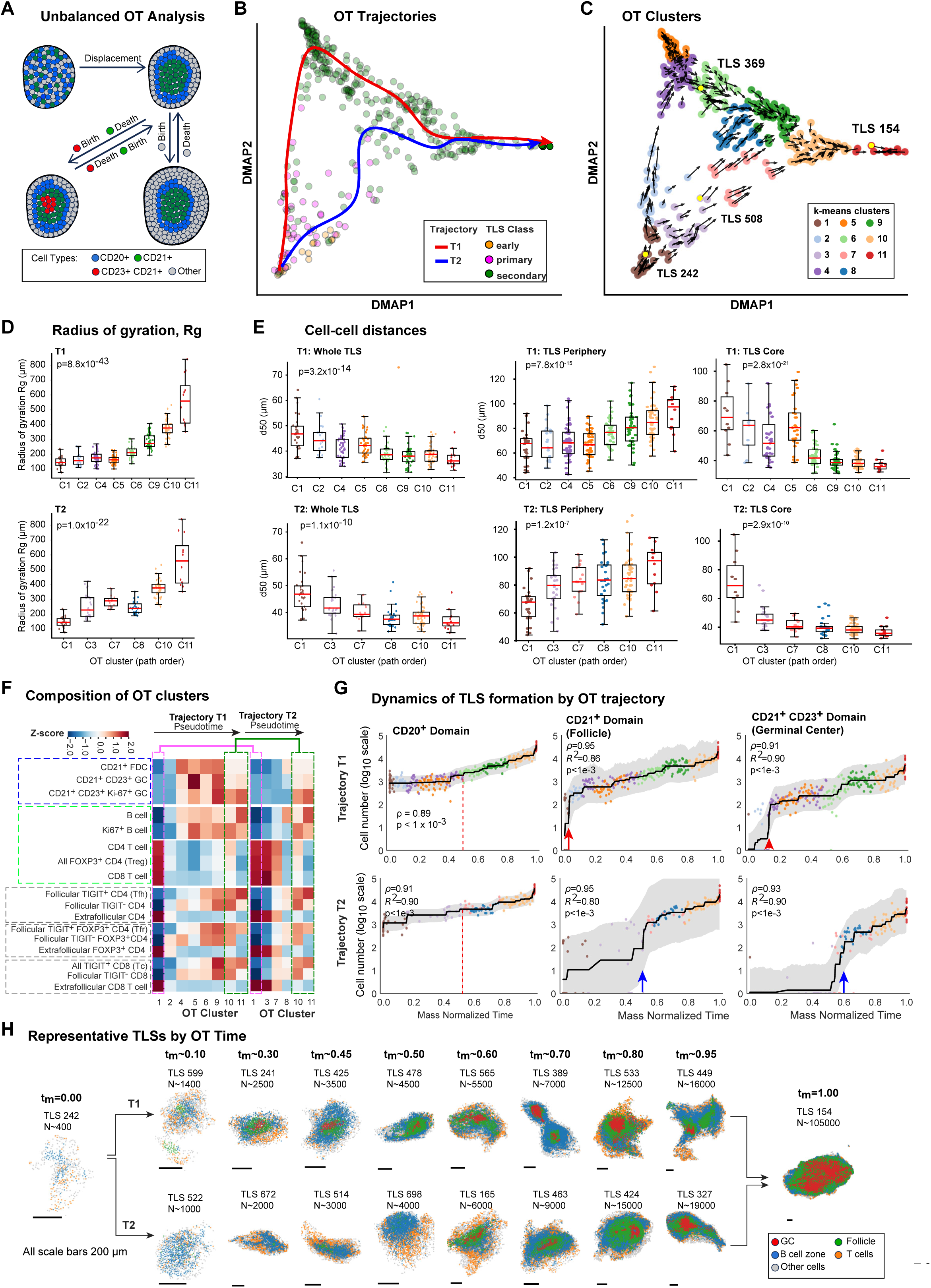
Optimal transport-based modeling reveals continuous and divergent TLS maturation trajectories. **(A)** Schematic of the optimal transport (OT) framework used to compute pairwise TLS distances. Each TLS is represented as a distribution of phenotyped single cells in a TLS-centered 3D space. Pairwise differences in spatial and phenotypic organization are quantified using an entropy-regularized, unbalanced fused Gromov–Wasserstein (UFGW) distance (see Supplemental Methods), capturing cell displacement, birth, and death events required to transform one TLS into another. Distances were computed across four phenotype groups: all cells, CD20⁺, CD20⁺CD21⁺, and CD20⁺CD21⁺CD23⁺. **(B)** Principal curves fitted to Trajectories T1 and T2 define pseudotime along each trajectory. TLSs are colored by early, primary, and secondary classification, illustrating continuous progression across the embedding rather than discrete stage boundaries. **(C)** Diffusion-map visualization of TLS maturation trajectories, with k-means clustering defining discrete regions along the manifold. Highlighted in yellow are TLS 242 (tₘ = 0.00), TLS 369 (tₘ = 0.52 on T1), TLS 508 (tₘ = 0.52 on T2), and TLS 154 (tₘ = 1.00). See also **Figure S6E**. **(D)** Radius of gyration increases along both trajectories, reflecting progressive TLS growth. **(E)** TLS compactness across trajectories and spatial compartments. Scatter box plots show the distribution of median k-nearest neighbor distances (d50, µm) for individual TLSs across ordered OT clusters along T1 (top) and T2 (bottom), including the whole TLS as well as periphery (0–10 µm from the TLS surface) and core (>20 µm from the TLS surface) regions. In all panels, boxplots represent the first and third quartiles with the median indicated by the central line, and individual TLSs are shown as overlaid points. *P* values shown following a Kruskal–Wallis test. **(F)** Heatmap of z-scored cell-type abundance across OT clusters along both OT trajectories highlighting differential enrichment patterns in B cell proliferation, follicular organization, and germinal center features along the two trajectories. **(G)** Trajectory-specific maturation dynamics of follicular zones along mass-normalized OT flow time (tₘ). CD21⁺ and CD23⁺ zones expand earlier along T1 than T2, indicating delayed follicular maturation along T2. Shaded bands show 95% bootstrap intervals. **(H)** Maximum projections of representative TLSs at matched tₘ along both paths, illustrating divergence in follicular organization. Cells are colored as CD20⁺, CD21⁺, CD23⁺, T cells, and “Other cells”). Scale bars are 200 µm.

OT reconstruction generated two overlapping trajectories for TLS assembly (T1 and T2), with the smallest and least mature TLSs located at the origin of the diffusion map (lower left in **Figure 4B**) and the largest and most mature TLSs at the upper right. TLSs classified as early or primary by the traditional EPS scheme mapped predominantly on the lower left and were substantially intermixed with each other, whereas TLSs classified as secondary were distributed across much of the diffusion map (**Figure 4B**). This finding underscores the extent to which the EPS scheme failed to account for the majority of TLS variation captured by the OT framework.

To determine whether reconstructed trajectories were physically and biologically plausible, we examined properties associated with TLS maturation. First, we applied *k*-means clustering to generate a set of 11 representative clusters: 8 in T1 and 6 in T2 with 3 common to both (**Figure 4C**). Along T1 and T2, successive clusters exhibited progressive and significant increases in the size of the TLS (as measured by radius of gyration, Rg; **Figure 4D**) and decreasing cell-to-cell distance (as measured by the median distance of each cell to 50 nearest neighbors; d50; **Figure 4E**). In zones proximal to TLS boundaries, d50 distances increased with OT cluster progression, indicating dispersion of cells away from the TLS boundary due to out-migration of lymphocytes. In contrast, within the core of each structure, defined broadly as the region >20 µm from the TLS boundary, d50 distances decreased monotonically with pseudotime (**Figure 4D**). These data are consistent with evidence that the follicular cores of TLSs become steadily more consolidated and organized as they mature,^68,69^ and shows that OT reconstruction captures these trends in a more precise manner than traditional EPS classification.

When we scored cell type enrichment along OT trajectories, we found that CD21⁺ FDCs were gradually replaced by CD21⁺CD23⁺ FDCs in GCs; these GCs commonly contained Ki67+ cells of multiple types, not all of which could be distinguished due to lateral spillover (blue dashed outline; **Figure 4F**). Lateral spillover occurs when segmentation errors cause antibody signals arising from densely packed or highly irregular cells to contaminate each other (lateral spillover can also be exploited to identify cells of different types in intimate contact).^70–72^ FDCs that are part of a GC are prototypical of cells that are difficult to segment. The number of B cells increased with cluster number, but overall T cell count fell (green outline). In contrast, T cell lineages associated with follicular and GC activity, such as T-follicular helper (Tfh) and regulatory (Tfr) cells, exhibited progressive increases. To compare trajectories T1 and T2 directly, we computed mass-normalized flow time (tₘ), which aligns TLSs with comparable numbers of cells in the two trajectories while preserving trajectory order (**Figure 4G**). Along T1, we observed coordinated formation and expansion of CD21⁺ FDC networks and CD23⁺ CD21⁺ GCs earlier and more rapidly relative to TLS size (**Figure 4G**; red arrows) than along T2 (blue arrows). The timing of FDC consolidation and follicle formation was also more heterogenous along T2 than T1 and GCs were generally smaller as a proportion of follicle (and TLS) volume along T2. These differences were evident in projections of representative TLSs aligned by tₘ (**Figure 4H**; see also **Figure S6E).** Here, GCs which are depicted as regions of contiguous mature FDCs (labelled in red), first appeared in trajectory T1 at tₘ= 0.32 (TLS 241) but only at tₘ= 0.63 (TLS 165) in T2, and were substantially smaller as a proportion of the follicular domain (represented by green label) and the entire TLS. Overall, these data show that both trajectories generated by OT corresponded to progressive changes in TLS architecture and biological activity, but that T1 and T2 differ in the (relative) time at which follicular consolidation and GC formation occur; below we show that TLS in T1 and T2 also differ in their physical environment.

### High resolution 3D confocal imaging reveals functional properties of maturing TLSs

A striking feature of reconstructed OT trajectories was the diversity of secondary TLSs; for example, structures with well-formed CD21^+^ CD23^+^ FDC networks were found at both the upper left (C4 to C6; **Figure 4C**) and the upper right (C10, C11) of the diffusion map landscape. To confirm that these TLSs had cell types and interactions typical of functional GCs, we performed high resolution CyCIF on sections ∼35 µm thick using a portion of the tissue block that immediately adjoined the serial section reconstruction. One TLS imaged at high resolution was structurally similar to clusters 4 to 6 along T1 with the follicle occupying much of the TLS; the second TLS was representative of structures in clusters 10 to 11 (**Figures 5A, 5B**). In TLS-A, CD21 and CD23-expressing FDCs formed a dense network (green) of interdigitating dendrites that surrounded B cells. This cage-like structure was itself surrounded by CD31⁺ αSMA⁺ blood vessels and by CD4 and CD8 T cells. A vessel positive for the endothelial marker CD31 and peripheral node addressin (PNAd; arrow) was observed to run through the center of the FDC network. This likely corresponded to a high-endothelial venule (HEV), a specialized post-capillary vessel found in lymphoid organs that acts as entry point for circulating lymphocytes. TLS-A was also characterized by physical interaction between PD1⁺ receptor on Tfh cells and PDL2 ligand expressed on CD21⁺ FDCs. Juxtracrine signaling between Tfh cells and FDCs is known to be mediated by PD1-PDL2 binding^73^ and functions in localized antigen presentation and B cell selection within the follicle.

**Figure 5.**
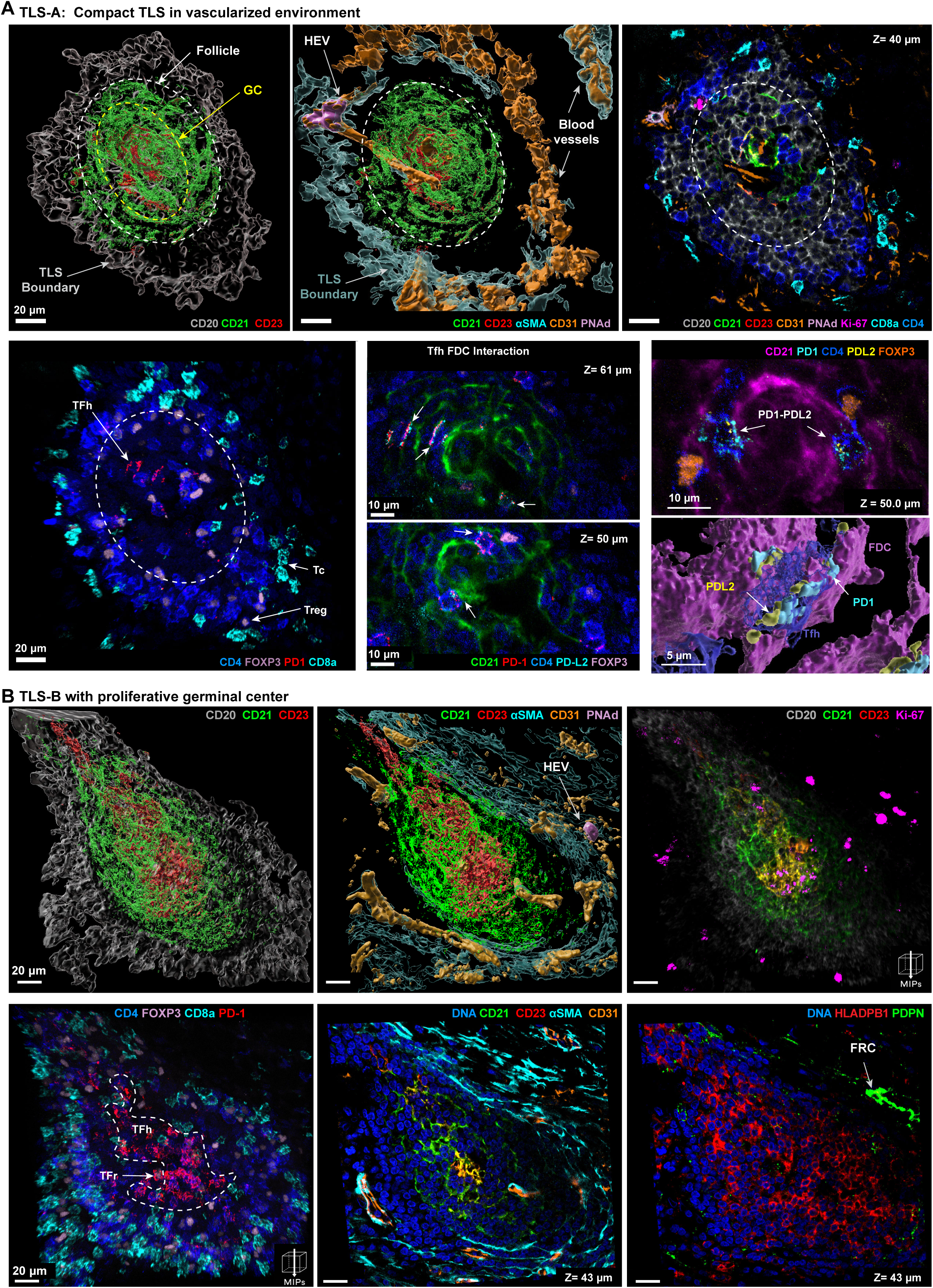
High-resolution 3D imaging reveals vascular integration and distinct follicular architectures within secondary TLSs. **(A)** Three-dimensional reconstruction of a TLS (TLS-A) embedded within vascular structures, showing an αSMA⁺CD31⁺ vessel extending into the follicle and PNAd⁺ high endothelial venules. Representative optical planes (280 nm spacing) illustrate compartmentalized CD20⁺ B cell zones, CD21⁺ follicle, and CD21⁺CD23⁺ FDC networks, with CD4⁺FOXP3⁺ Tregs localized within follicles, CD8⁺ cytotoxic T cells at the periphery. Also shown are PD-1⁺ Tfh cells interacting with PD-L2⁺CD21⁺ FDC dendrites. **(B)** Thick-section confocal imaging of a mature TLS (TLS-B) demonstrating a highly proliferative germinal center. Surface renderings highlight CD21⁺CD23⁺ follicular dendritic cell (FDC) networks with an architecture distinct from that observed in TLS-A. Maximum-intensity projections show Ki-67⁺ proliferating B cells, PD-1⁺ Tfh cells, PDPN⁺ fibroblastic reticular stromal networks, and HLA-DPB1⁺ antigen-presenting cells. Scale bars, as indicated.

The second TLS (TLS-B) also had dense CD21⁺ CD23⁺ FDC networks but the morphology of the dendrites was different, and the TLS contained more Ki-67⁺ B and Tfh cells both within and outside of the follicle (dashed outline). A probable HEV was present (arrow), along with vascularization as in TLS-A. Other relevant features included PDL2^+^ FDCs in association with PD1^+^ Tfh, networks of cells positive for podoplanin (PDPN), a mucin-type glycoprotein that is a marker of fibroblastic reticular cells (FRCs), and expression of MHC class II molecules (HLA-DPB1) on B cells (**Figure 5B**). Thus, both TLS-A and TLS-B have multiple features associated with advanced follicular specialization including cell types and cell-cell interactions associated with antigen presentation, but within the context of distinct 3D structures.

### TLS maturation trajectories correspond to increasing GC activity

To further quantify the link between OT trajectories and establishment of a functional GC microenvironment, we measured association of B and Tfh cells with FDCs based on patterns of CD21, CD86, and CXCL13 staining (this analysis used the entirety of the standard-resolution CyCIF data, not just 3D data). CD86 is a costimulatory ligand for T cell activation that is expressed by several types of antigen presenting cells (APCs) including B cells.^74–77^ CXCL13 is a chemokine expressed by FDCs, Tfh cells, and some stromal cells that functions as a chemoattractant for CXCR5-expressing B and T cells. CXCL13 levels are an established measure of lymph node GC activity, for example, following vaccination.^78^

We observed that CD86 primarily stained CD20^+^ B cells in GCs and follicles (labelled “1” in **Figures 6A-B);** many of which were in the proximity of FDCs (“2”); Tfr and Tfh cells were also found nearby (“3 & 4”). In contrast, CXCL13 staining had two patterns: (1) punctate staining of FDCs and Tfh cells (“5 &6”) and (2) extended staining of fibers or vessels (“7”) that did not correspond to CD31^+^ blood vessels (“8”). 3D high resolution imaging showed that the punctate pattern was consistent with secretory granules and the second pattern (which was ∼6-folder weaker in maximum intensity) corresponded to reticular structures with a narrow lumen formed from cells with spindle-like nuclei (**Figure 6C, S6F)**. Similar structures are discernable in published images of CXCL13 staining in lung cancer specimens.^44^ We interpret them to represent lymphatic vessels (LVs; although confirmatory staining for the lymphatic endothelial cell marker LYVE-1 has not yet been successful in any of our 3D experiments). Both lymphatic endothelial cells and associated stromal cells have been reported to express CXCL13 under some circumstances^79^. However, it is also possible that the pattern we observe arises from binding of CXCL13 to heparin-sulfate ECM on LVs. In this interpretation, the two patterns of staining we observe correspond to the two components of soluble and immobilized CXCL13 gradient.^80^

**Figure 6.**
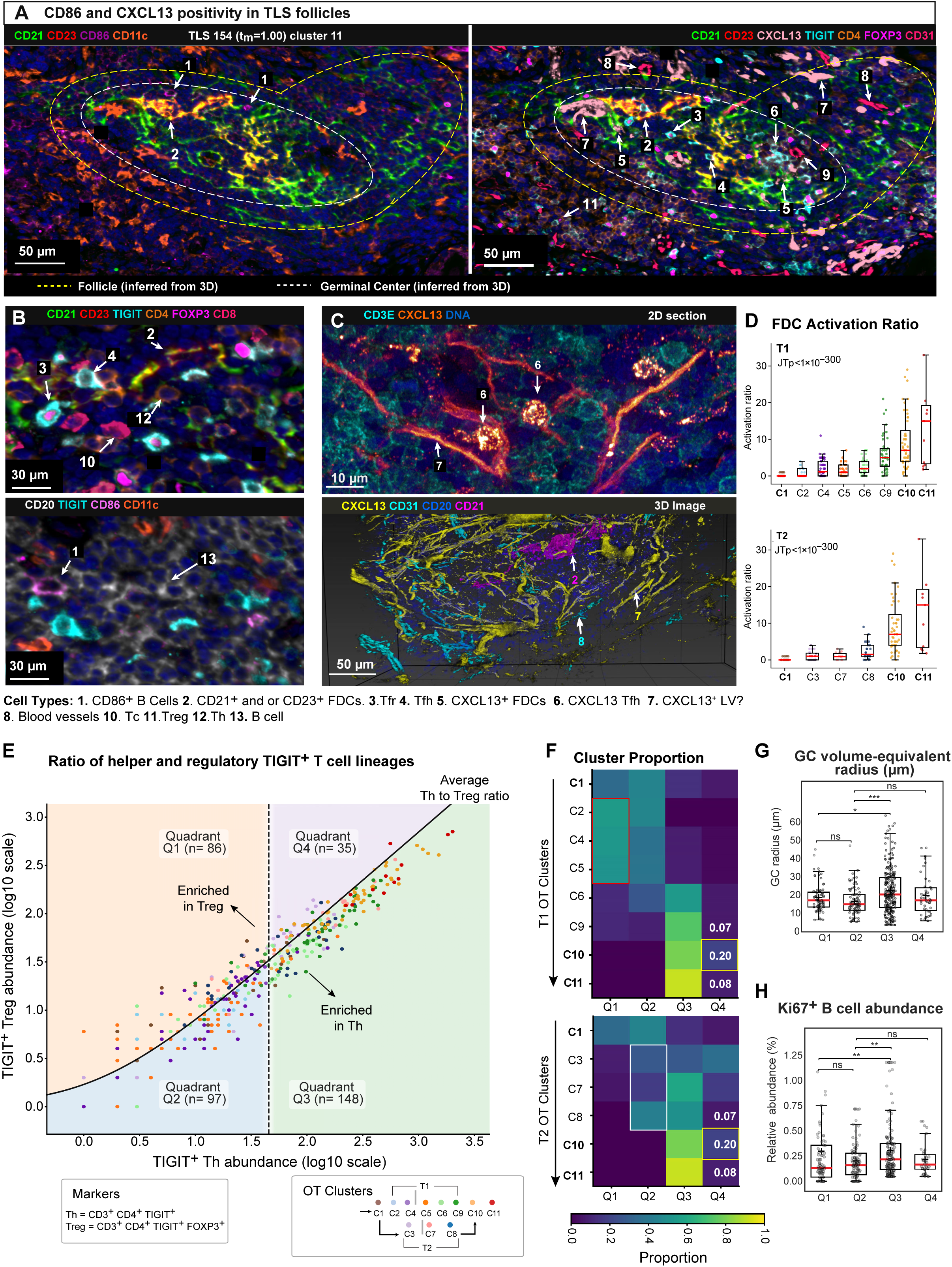
Optimal transport-defined TLS trajectories reveal progressive follicular activation and divergent immune regulatory states. **(A)** Representative image cross-sections of a mature TLS (TLS154, cluster 11) showing CXCL13⁺ and CD86⁺ regions in TLS follicle proximity to TIGIT⁺ T cells. Numbered regions highlight areas of interest **(B)** Corresponding high-magnification insets from A (bottom panels). **(C)** Representative high-resolution confocal image (top panel) showing CXCL13-expressing lymphatic vessel adjacent to T cells with punctate CXCL13 expression. Bottom panel shows CXCL13-expressing lymphatic vessels adjacent to CD31⁺ blood vessels and CD21⁺ follicular dendritic cell networks within TLS-associated tissue. **(D)** Activated FDC ratios increase across clusters along both paths. Ratios are defined as CD86⁺CXCL13⁺CD21⁺ FDCs divided by 1+ (CD86⁻CXCL13⁻CD21⁺ FDCs). **(E)** Joint analysis of TIGIT⁺ CD4 T cells and TIGIT⁺ Tregs stratifies TLSs into four immunoregulatory quadrants (Q1-Q4) based on abundance and relative balance. Points represent individual TLSs colored by OT cluster. The vertical dashed line indicates the median TIGIT⁺ Th abundance, and the diagonal line indicates the average TIGIT⁺ Treg:Th ratio (see Methods). **(F)** Heatmap showing the distribution of TLSs across Q1-Q4 quadrants by OT cluster, showing shifts in immunoregulatory states along both trajectories. **(G)** Germinal center size varies by quadrant, with larger volume-equivalent radii in the high-Th cell, high-Tfh enrichment quadrant (Q3), compared to the low-Th cell quadrants (Q1 and Q2). **(H)** Proliferative activity of B cells across quadrants. Shown is the relative abundance of Ki-67⁺ B cells normalized to total cells within the combined GC, CD21⁺, and CD20⁺ compartments of each TLS.

Given the complexity of these expression patterns, we again used a microregional labeling strategy in which the fraction of CD21^+^ FDCs with CXCL13 and CD86 staining within the same segmentation mask (∼10 µm diameter) or not, was used to compute an “activation ratio”. We observed steady and significant increases in this ratio along trajectories T1 and T2 with the strongest increase observed in the most advanced TLS clusters (**Figure 6D**). Similar calculations were performed for CD86-FDC association alone, with a range of different segmentation approaches, to focus on punctate (secretory granule) versus diffuse (lymphatic) CXCL13 staining, with similar overall results (**Figure S6G**). We conclude that both trajectories inferred by OT correspond to an increasingly functional GC microenvironment characterized by CXCL13-mediated lymphocyte recruitment.

The activity of germinal centers is regulated in part by the relative abundance of Tfh and Tfr cells^81,82^ and their ratio is a commonly measured blood diagnostic.^83^ The ratio in our dataset averaged ∼3 across secondary TLS GCs but with a high level of variance (**Figure S5E**). To study how the helper-to-regulatory T cell proportion changed along OT trajectories, we compared the prevalence of all TIGIT^+^ Th and Treg cells in the TLSs without regard to zonation, allowing comparison of early, primary and secondary TLSs. We then divided the landscape into four quadrants (Q1-Q4) based on (i) overall TIGIT^+^ Th cell prevalence (X axis) and (ii) the TIGIT^+^ Treg to Th ratio (Y axis; each point represents one TLS, color-coded by OT trajectory; **Figure 6E**). At earlier tₘ values (C1 to C5), TLSs lay mostly in Q1 and Q2. T1 clusters were slightly enriched in Tregs (red outline; **Figure 6F**) and T2 in Th cells (white outline). At later tₘ values (C6 to C11), a shift towards Q3 was observed, reflecting an increase in the Th to Treg ratio. This was accompanied by significant increases in GC volume (**Figure 6G**) and the proportion of proliferating B cells (**Figure 6H**). The progressive nature of these architectural and proliferative changes further supports the interpretation that OT trajectories primarily represent paths of increasing TLS maturity and activity.

OT cluster 10 was notable however, in exhibiting a 3-fold increase in Q4 (more repressed) TLSs as compared to either C9 in trajectory T1 or C8 in trajectory 2 (yellow outlines; **Figure 6F**). It is possible that this increase reflects Treg-mediated attenuation of germinal center responses.^84^ The data we used for OT analysis cannot disentangle TLS regression from TLS growth, but monotonic increases in nearly all of the physical and functional properties of TLS maturity analyzed thus far suggest that regression is unlikely to be a dominant feature of the data, except perhaps for TLSs in C10. Future 3D OT analysis of longitudinal data, from mice for example, may improve our ability to distinguish TLS regression from growth.

### Distinct three-dimensional spatial localization of TLS maturation states in the tuberculosis lung

To determine if proximity to granulomas impacts TLS maturation, we identified GrALTs as TLSs located within 100 µm of a granuloma in any single tissue section. Overall, 62% of TLSs were GrALTs (**Figures S4A**) and there were significantly enriched for trajectory T2 (79%) versus T1 (59%) (Fisher’s exact test, p = 8.11 × 10⁻^6^; **Figures 7A-B**). To identify tissue features associated with T1, we used spatial latent Dirichlet allocation (LDA) to identify TLS-associated recurrent cellular neighborhoods (RCNs; **Figures S7A-C**). We found that TLSs in T1 clusters 2, 4, 5 were preferentially proximal to RCN3, which corresponded to a CD31⁺ αSMA⁺ perivascular neighborhood (**Figures 7C, S7C-D**). High resolution images of TLS-A provided a striking example of a TLS surrounded by multiple blood vessels (**Figure 5A**). We conclude that the TLSs in T2 are preferentially granuloma associated, suggesting a role for the GME in TLS maturation, whereas compact TLSs in T1 are associated with a rich vasculature.

**Figure 7.**
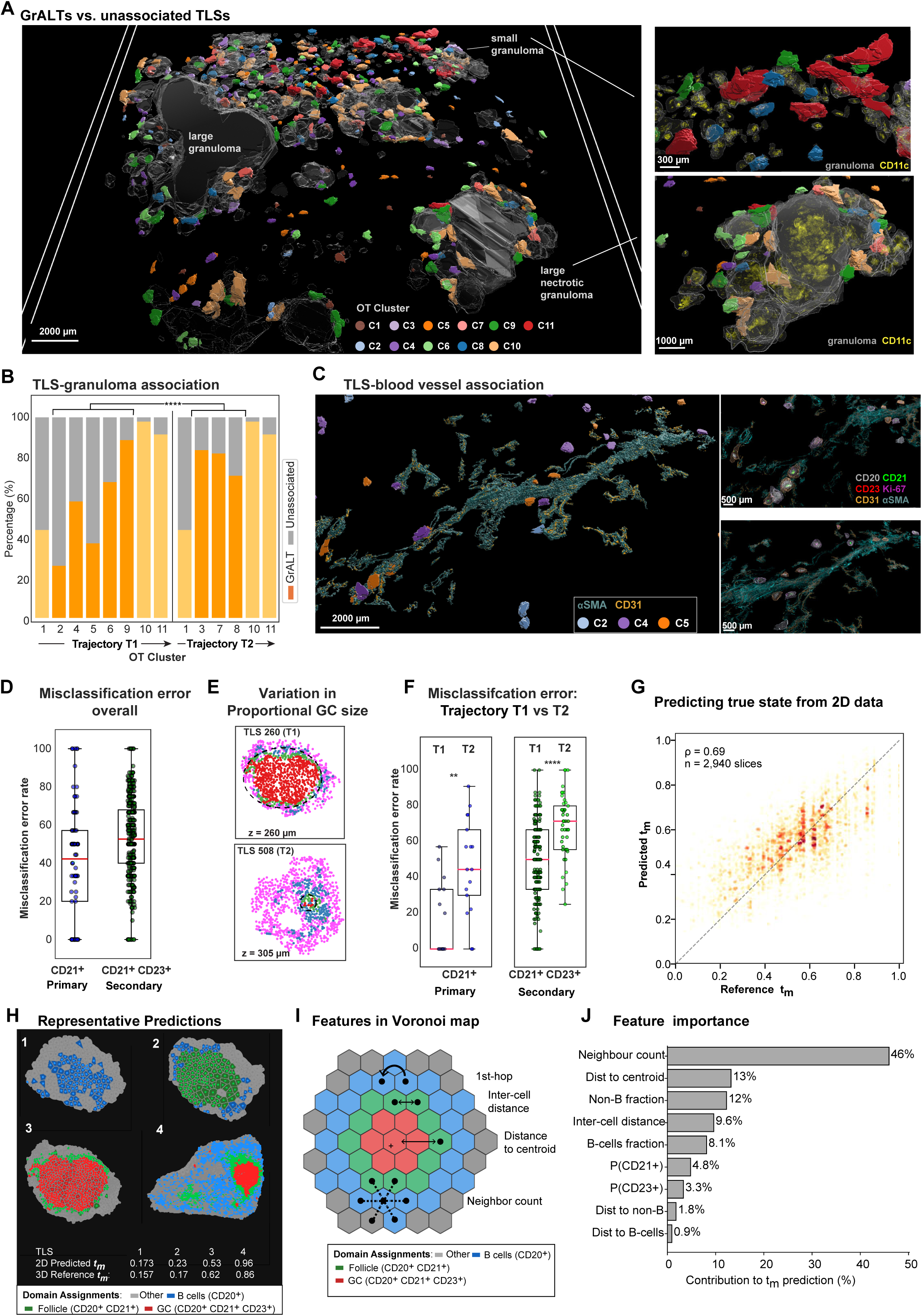
Spatial organization of TLS maturation states and resolution of sectioning-dependent misclassification in pulmonary TB. **(A)** 3D rendering of TLSs within reconstructed lung tissue, colored by OT cluster and shown relative to granulomas (gray), illustrating both granuloma-associated and unassociated TLSs. Representative TLSs are adjacent to small granulomas and large necrotic granulomas associated with multiple TLS clusters. **(B)** Proportion of granuloma-associated lymphoid tissues (GrALTs) and unassociated TLSs across OT-defined clusters in T1 and T2. **(C)** Representative TLSs from T1 clusters positioned adjacent to CD31⁺αSMA⁺ vascular structures. **(D)** Misclassification error rates for primary and secondary TLSs inferred from single 2D sections, highlighting the limitations of 2D TLS staging. **(E)** Representative 2D sections illustrating differences in the relative sizes of follicle and GC compartments between in representative T1 and T2 TLSs. **(F)** Misclassification rates stratified by OT-defined trajectories (T1 versus T2). Boxplots show median and interquartile range; points represent individual TLSs. Statistical significance by Mann-Whitney U test; P < 0.01 (**), P < 0.0001 (****). **(G)** Prediction of 3D TLS maturity (tₘ; mass-normalized time) from single 2D TLS sections. Predicted versus true 3D tₘ (ρ = 0.688, n = 2,940; color indicates point density). **(H)** Representative Voronoi phenotype maps of 2D TLS slices across predicted TLS maturity states, with cells colored by phenotype. Predicted 2D and reference 3D tₘ scores are indicated. Examples illustrate how individual sections from large or mature TLSs can variably resemble earlier-stage TLSs, contributing to reduced prediction accuracy at high 3D tₘ values. See also **Figure S7G**. **(I)** Spatial features used for prediction of 3D TLS maturity from 2D sections. Schematic illustrating representative graph-derived spatial features extracted from Voronoi-based cellular neighborhoods, including neighborhood (hop-based) connectivity, inter-cell distance, distance to local centroid, and neighbor count across CD20⁺, CD20⁺CD21⁺, CD20⁺CD21⁺CD23⁺, and non-follicular non-B cell regions. **(J)** Relative contribution of feature categories to the prediction of 3D tₘ in the gradient boosting model. Neighborhood connectivity and centroid-distance features contributed most strongly to maturity prediction, indicating that higher-order spatial organization and compartmentalization encode substantial information about 3D TLS state in individual 2D sections.

### Comparing TLS staging by 2D and 3D imaging

Finally, we re-examined misclassification of TLS stage (by EPS criteria) when only a single 2D tissue section was used to assess CD21 and CD23 positivity, using 3D imaging data as the ground truth (**Figures 7D, S7E**). We found that primary TLSs were misclassified 41% of the time on average and secondary TLSs 53% of the time (**Figure 7D**). Simple geometric considerations suggest that misclassification rates should depend on the fraction of TLS volume encompassed by FDCs: the larger the relative size of the germinal center the harder it is to miss in 2D. This likely explains why TLSs associated with OT trajectory T2 were more frequently misclassified than those in T1 (**Figures 7E-F)**. Moreover, in tonsil, GCs often occupy a much larger fraction of the structure (up 70% of the volume) than in a TLS (∼3% of volume), potentially explaining why high rates of 2D misclassification have not previously been described.

To determine whether computational models can infer the true maturation state of a 3D TLS from either a single or a limited number of 2D slices from the same TLS, we trained histogram-based gradient boosting regressors using spatial summary features. The mass-normalized flow time (tₘ) of the TLS, as estimated by OT from 3D data, was used as the reference for TLS maturity state (**Figures 7G-H, S7F**). First, we confirmed that the model was able to correctly predict the reference tₘ when consecutive slices were omitted (Spearman ρ ≈ 0.96; R² ≈ 0.92), with performance improving steadily as additional sections were incorporated (**Figure S7F**). Remarkably, even a single 2D section retained substantial predictive signal for tₘ (ρ ≈ 0.69) and recapitulated progressive enrichment of B cell, follicular, and GC zones in representative sections (**Figures 7G-H**). Prediction accuracy was highest for TLSs with intermediate tₘ values (0.25–0.75), possibly because marginal sections from the most mature TLSs resemble earlier-stage TLSs, confusing the model (**Figures S7G, 7H**). Overall, we conclude that, when interpreted within an appropriate computational framework, 2D sections retain substantial information about 3D TLS organization and outperform conventional EPS classification (**Figures 7D, 7F**).

What properties of 2D images make them predictive of a 3D structure? Feature-importance analysis showed that predictive features included cellular composition (the proportion of non-B cells), neighborhood structure, TLS geometry and degree of compaction (this corresponded to model parameters neighbor count, distance to TLS centroid, inter-cell distance etc.; **Figures 7I-J**). Thus, the organization and composition of cells in a 2D section of a TLS are measurably influenced by the presence of a follicle or GC even when that structure is not within the same tissue section. Further development of models for 2D to 3D inference should therefore be valuable for the many studies in which 3D reconstruction is not feasible.

## DISCUSSION

The current model of TLS development, which is based on traditional histology, flow cytometry, scRNAseq, and 2D spatial transcriptomics^85,86^ postulates sequential TLS assembly via focal recruitment of B and T cells to create an early (E) TLS, formation of a follicular dendritic network in a primary (P) TLS, and maturation of the FDC network in a secondary (S) TLS to create a functional GC. In this EPS maturation trajectory,^87^ the most substantial morphological and structural changes occur at early to primary and primary to secondary transitions. However, 3D reconstruction of hundreds of TLSs in *Mtb*-infected lungs shows that EPS classification based on 2D imaging does not accurately capture TLS maturation state, leading to misclassification rates of 40% or more. It also misses the fact that up to 20% of TLSs are multi-focal (i.e., have at least two independent FDC foci of GCs). The primary consequence of this is an overestimation of the number of early and primary TLSs and an underestimation of secondary TLSs, the maturation stage most closely associated with functional immune responses. 2D imaging also fails to capture the wide diversity of secondary morphologies, which vary several hundred-fold in cell number, can have multiple germinal centers, and exhibit more or less separation between B and T cell-rich zones (zonation).^88^

### Reframing TLS development using optimal transport

We find that the diversity of secondary TLSs can be organized in a logical and biologically plausible manner using optimal transport. This is a form of trajectory analysis in which structures are placed along a pseudotime trajectory based on size and morphology, as determined by minimizing the number of cell birth, death, and transport events needed to make successive structures maximally similar. Trajectory analysis is common in dissociative and spatial scRNA analysis^89^ and OT is widely used to compare 3D structures in engineering and material science; it has also been applied to volumetric radiological data^90^ but not, to our knowledge, to 3D tissue structures. OT trajectories are commonly interpreted as developmental trajectories, and this makes particular sense in the current setting because OT approximates the physical process by which TLSs form via accumulation of cells from the circulation, proliferation within the TLS, and out-migration as immune cells mature. These fluxes generate interconnections among TLS zones, likely explaining why TLSs, follicles, and GCs exhibit allometric scaling relationships

Modeling with OT confirms that early and primary TLSs are more similar to each other than secondary TLSs architecturally, and that secondary TLSs form several distinct classes. This diversity gives rise to two developmental trajectories. In the first trajectory (T1), FDC networks and germinal centers form when the overall structure is still compact leading to assemblies in which the size (radius of gyration) of the FDC network is a substantial portion of the B and T cell aggregate. In the second trajectory (T2), FDC consolidation and germinal center formation is delayed until the structure is ∼10-fold larger in size. Several properties of these trajectories suggest that they represent a real biological series. First, both are characterized by an overall increase in the size of the B cell compartment, tighter cell packing specifically in the core of each structure and decreasing proportions of non-follicular stromal and myeloid cells. The inverse relationship between TLS size and median cell-cell distance is consistent with chemokine-guided follicular organization and consolidation and with germinal center-associated antigen selection.^74,75^ Second, markers of biological activity such as B cell proliferation, maturation of FDCs, recruitment of Tfh cells, and regional expression of CXCL13 all increase monotonically along OT trajectories. Finally, high resolution thick-section CyCIF of selected TLSs demonstrates the presence of interactions between B and T cells and FDCs associated with active immune programing, including PD1⁺ Tfh cells in direct contact with PDL2⁺ CD21⁺ FDC networks. We also observed other features including PDPN⁺ fibroblastic reticular cell networks. We conclude that trajectories inferred by OT accurately capture key features of TLS biology.

### Association of TLS morphology with other tissue structures

We find that TLSs identified by OT to be the most mature (i.e., where T1 and T2 converge) are almost always (>90%) associated with granulomas. This observation is consistent with prior studies showing that TB granulomas are antigen- and cytokine-rich microenvironments that induce and organize peripheral lymphoid tissue.^91–93^ As more and larger 3D datasets become available, it will be possible to more precisely determine how TLS maturation relates to granuloma type (necrotizing versus non-necrotizing) and disease stage. In contrast to large fully developed TLSs, less mature clusters along trajectory T1 were frequently proximal to networks of blood vessels. These vessels are in addition to PNAd⁺ vessels that penetrate the follicles and germinal centers (and likely represent HEVs). In this context, it is noteworthy that granulomas are themselves 3D structures with a characteristic ginger root morphology^94^ at a mm to cm scale, 10-100 fold larger than that of a TLS. Accurately mapping TLS and granuloma associations will likely require integration of X-ray microtomography and fluorescence tissue imaging. In the context of *Mtb*, the 3D co-evolution of TLSs and granulomas may provide important insights into bacterial control.

Nearly all of TLSs described in this study are associated either with bronchioles or what we interpret to be the remnants of bronchial epithelium and are therefore likely to represent induced bronchus-associated lymphoid tissues (iBALTs).^95^ This presumably reflects the response to persistent *Mtb* bacilli. We do not yet know whether TLSs present in post-TB disease, where fibrosis is much more extensive, or other lung diseases will be similar to the ones described here. However, pilot studies in non-small cell lung cancer and colorectal cancer show that 2D misclassification of TLS maturation state will be an issue outside of the setting of pulmonary TB (**Figures S2A-B**). Collecting additional 3D data on TLSs will benefit from new imaging approaches, such as cyclic light sheet fluorescence microscopy,^96^ that reduce the cost and high level of effort and associated with multiplexed serial section reconstruction.

### Implications for studying TLS biology

Our data may call into question some studies that examine the association between TLS maturity and disease progression or response to therapy based on 2D datasets alone:^97^ it is likely that some 2D studies have missed key distinctions between TLS stages or immunoprotective versus immunosuppressive biology. Additional 3D data from a variety of diseases will help to address this concern but it is inevitable that most studies of tissue resident immunity will continue to rely on 2D sections: routine 3D imaging of large human cohorts is too time-consuming. It is therefore important to understand what can and cannot be learned from 2D imaging. When a 2D section successfully intersects a GC core, then maturation state can be reliably assessed. However, the absence of CD21⁺CD23⁺ FDC networks in a 2D image is not a definitive indication of the absence of a GC. Thus, 2D imaging represents a high precision low-recall assessment of true TLS state. The extent to which this impacts biological interpretation will vary with follicle morphology: the larger the germinal center as a proportion of the overall assembly the less likely it is to be missed. In tonsil, for example, germinal centers are large compared to follicles, and misassignment of maturation state is relatively infrequent. Similarly, TLSs on trajectory T1 were less likely to be misclassified than those on T2, simply because the former were more compact.

We provide evidence that 2D images contain information on 3D states that are not visible by simple inspection of CD20, CD21, and CD23 staining patterns. Using down-sampling approaches and CNN models, we find that 2D sections are likely to contain substantial “hidden” information on 3D organization. As a result, TLS maturity predicted from single 2D sections correlates with the ground truth 3D OT development stage (pseudotime value) with Spearman ρ ≈ 0.69. Computational analysis aimed at identifying the precise nature of this information will be an important step in reliably linking 2D and 3D datasets.

### Limitations and Practical Implications

The current study has several limitations, most notably the limited number of disease stages in which TLSs have been reconstructed. With the availability of additional data, the OT trajectories identified in this paper are likely to change or become more complex. Follow-up 3D studies involving additional specimens and larger Z dimensions will also be required to better map TLS association with granulomas. Moreover, spatial transcriptomics is substantially more effective than fluorescence imaging at detecting cytokines, which play a key role in TLS formation (**Figure S1C, S1E-G**), suggesting that volumetric image reconstruction should be paired with additional transcriptomic profiling – although this remains an expensive proposition. However, these concerns do not detract from the finding that computational methods such as OT can effectively organize key features of TLS immunology in 3D and thereby reveal hidden aspects of TLS maturation.

## STAR★ METHODS

### KEY RESOURCES TABLE

| REAGENT or RESOURCE | SOURCE | IDENTIFIER |
| --- | --- | --- |
| **Antibodies** |  |  |
| Mycobacterium tuberculosis Ag85B | Abcam | RRID: AB_776575 |
| CD86 | R&D Systems | RRID: AB_354385 |
| CD25 | LifeSpan BioSciences | RRID: AB_2889341 |
| Goat anti-Rabbit IgG-AF488 | Invitrogen | RRID: AB_2534114 |
| Donkey anti-Goat IgG-AF555 | Thermo Fisher Scientific | RRID: AB_2535853 |
| Goat anti-Mouse IgG-AF647 | Thermo Fisher Scientific | RRID: AB_2535806 |
| Granzyme B | Dako | RRID: AB_2114697 |
| CD3E-AF555 | Cell Signaling Technology | RRID: AB_3668891 |
| CXCL13 | R&D Systems | RRID: AB_2943272 |
| Goat Anti-Mouse IgG2a-AF488 | Jackson ImmunoResearch, Inc. | RRID: AB_2338855 |
| Goat Anti-Mouse IgG1-AF647 | Jackson ImmunoResearch, Inc. | RRID: AB_2338916 |
| PNAd | Sigma-Aldrich | RRID: AB_2943331 |
| FOXP3-eF570 | eBioscience | RRID: AB_2573609 |
| CD21-AF647 | Abcam | RRID: AB_2889188 |
| Goat anti-Rat IgM-AF488 | Thermo Fisher Scientific | RRID: AB_2535798 |
| CD23-AF488 | Abcam | RRID: AB_2943215 |
| Ki-67-PE | Cell Signaling Technology | RRID: AB_2797836 |
| CD20-eF660 | eBioscience | RRID: AB_11151691 |
| Cytokeratin (pan)-AF750 | Novus Biologicals | RRID: AB_2868569 |
| CD8a-AF488 | Thermo Fisher Scientific | RRID: AB_2574412 |
| TIGIT-AF555 | Cell Signaling Technology | RRID: AB_3674649 |
| PD-1-AF647 | Abcam | RRID: AB_2728811 |
| Alpha-actin-2-AF750 | R&D Systems | RRID: AB_2868436 |
| CD4-AF488 | R&D Systems | RRID: AB_2728839 |
| CD11c-AF555 | Cell Signaling Technology | RRID: AB_3331656 |
| PD-L1-AF647 | Cell Signaling Technology | RRID: AB_3669071 |
| PCNA-AF750 | Cell Signaling Technology | RRID: AB_3669072 |
| CD163-AF488 | Cell Signaling Technology | RRID: AB_3720265 |
| CD36-AF555 | Cell Signaling Technology | Cat# 40462BC; RRID: N/A |
| CD31-AF647 | Abcam | RRID: AB_2857973 |
| Mast Cell Tryptase-AF790 | Santa Cruz Biotechnology | RRID: AB_2943323 |
| CD11b-AF488 | Thermo Fisher Scientific (eBioscience) | RRID: AB_2637195 |
| CD206-AF555 | Cell Signaling Technology | RRID: AB_3674647 |
| CD45-AF647 | BioLegend | RRID: AB_493034 |
| Myeloperoxidase-AF790 | Santa Cruz Biotechnology | RRID: AB_2943295 |
| TCF1/TCF7-AF488 | Cell Signaling Technology | RRID: AB_2797627 |
| ICOS-AF555 | Cell Signaling Technology | RRID: AB_2800142 |
| IRF4-AF647 | Cell Signaling Technology | RRID: AB_2868605 |
| HLA-DRB1-AF488 | Abcam | RRID: AB_2889845 |
| CD68-PE | Cell Signaling Technology | RRID: AB_2799935 |
| CTLA4-AF647 | Abcam | RRID: AB_2943179 |
| Lamin-B1-AF488 | Abcam | RRID: AB_2728786 |
| Lamin-A/C | Cell Signaling Technology | RRID: AB_10997529 |
| NGFR-AF647 | BD Biosciences | RRID: AB_1645403 |
| ABCG1 | Novus Biologicals | RRID: AB_3188331 |
| **Biological samples** |  |  |
| FFPE tissue block (Patient1_B4) | Department of Pathology, Massachusetts General Hospital, Mass General Brigham | N/A |
| FFPE tissue block (Patient1_B5) | Department of Pathology, Brigham and Women's Hospital, Mass General Brigham | N/A |
| FFPE tissue block (Patient2) | Department of Pathology, Massachusetts General Hospital, Mass General Brigham | N/A |
| FFPE tissue block (Patient3) | Department of Pathology, Massachusetts General Hospital, Mass General Brigham | N/A |
| **Software and algorithms** |  |  |
| MCMICRO pipeline | https://doi.org/10.1038/s41592-021-01308-y | https://github.com/labsyspharm/mcmicro |
| SCIMAP (2.0.5) | https://doi.org/10.21105/joss.06604 | https://scimap.xyz |
| Adobe Illustrator (29.3.1) | Adobe | RRID:SCR_010279 |
| ImageJ (1.54p) | https://doi.org/10.1038/nmeth.2019 | https://imagej.nih.gov/ij/ |
| MATLAB 2022a | Mathworks Inc. | https://www.mathworks.com/products/matlab.html |
| Imaris 10.2.0 | Oxford Instruments; formerly Bitplane | [imaris.oxinst.com](https://www.oxinst.com) |

### LEAD CONTACT

Requests for further information should be directed to lead contact, Peter Sorger.

### MATERIALS AVAILABILITY

This study contains no unique reagents or resources; all antibodies are available commercially (see **Table S3** and key resources table)

### DATA AND CODE AVAILABILITY

Codes generated during this study will be available at the time of publication (https://github.com/labsyspharm/TB_3D_TLS_2026/). Image data generated during this study will be available via AWS transfer at the time of publication. Instructions for accessing the primary and derived data are available on a data index page via Zenodo (https://zenodo.org/10.5281/zenodo.19616347). GeoMx data are available as Count Matrix in **Table S1**. Raw files for GeoMx were submitted to Gene Expression Omnibus (Accession number: GSE318959).

### METHOD DETAILS

#### Tissue samples

Formalin-fixed paraffin-embedded (FFPE) tissue samples were obtained from biopsies and autopsies archived at Brigham and Women’s Hospital and Massachusetts General Hospital in Boston, Massachusetts (United States), and Africa Health Research Institute, University of KwaZulu-Natal, Durban (South Africa). Tissues were screened for *Mycobacterium tuberculosis* using acid-fast staining. Specimens were collected under MGH IRB 2019P001114, BWH IRB 2018P001627, and the University of KwaZulu-Natal Biomedical Research and Ethics Committee (BREC) protocol BE019/13.

Primary samples were lung biopsy from Patient 1 (50-year-old male, untreated, left upper lobe infiltrate with necrotizing and non-necrotizing granulomas). Corresponding 5-µm tissue sections were cut from sample blocks for subsequent processing. For the serial section reconstruction experiment, 120 adjacent sections were generated from the ∼2 cm x 1.5 cm x 600 µm Patient1_B4 tissue block, as outlined in **Figure 1; Table S2**. 40 H&E and 40 CyCIF images were obtained by skipping one intervening section between imaged sections, thereby capturing greater tissue depth which spanned ∼600 µm. Histopathological review and annotation of a subset of the H&E-stained serial sections was performed by a board-certified pathologist. Separately, 10 serial sections were cut from Patient1_B5 block; and Patient1_B5_Section 2 was processed for spatial transcriptomics using GeoMx Digital Spatial Profiler (DSP) platform, along with samples from Patient2 and Patient3 (Patient2: a 64-year-old male with COPD and alcoholic cirrhosis with a surgical lung biopsy for a 1.6 cm lung nodule; Patient3: a 62-year-old female with rheumatoid arthritis who was PPD-positive and received 5 months of isoniazid therapy for latent TB). Complementary 35-µm thick sections from Patient1_B4 were analyzed by high-resolution multiplexed confocal imaging (3D CyCIF),^98^ enabling detailed spatial characterization of TLS architecture and cellular interactions that validated features observed in whole-slide 3D reconstruction.

#### 2D image acquisition (H&E and tissue-based CyCIF protocol)

H&E-stained FFPE sections were scanned on an Olympus VS200 automated slide scanner equipped with a 20X objective (NA 0.8; 0.2738 µm/pixel) at the Neurobiology Imaging Facility at Harvard Medical School. Multiplexed CyCIF imaging was performed on FFPE tissue sections as previously described.^99,100^ A detailed reproducible protocol is available at protocols.io (https://dx.doi.org/10.17504/protocols.io.5qpvorbndv4o/v2). Slide preprocessing was carried out using a BOND RX Automated IHC/ISH Stainer. Sections were initially baked at 60°C for 30 minutes and subsequently dewaxed with the Bond Dewax reagent at 72°C. Heat-induced epitope retrieval was performed using Leica Epitope Retrieval 1 solution at 100 °C for 20 minutes. Slides were then processed through multiple iterative rounds of antibody staining, imaging, and fluorophore bleaching. The complete list of primary and secondary antibodies used in the serial section reconstruction CyCIF experiment is provided in **Table S3**. Antibody incubations were conducted overnight at 4 °C in the dark, and Hoechst 33342 was included in each staining cycle to label nuclei. After washing steps, coverslips were mounted in 50% glycerol prepared in PBS. Fluorescence images were captured on a CyteFinder slide-scanning microscope (RareCyte Inc. Seattle WA) equipped with a 20×/0.75 NA objective. Fluorophore inactivation between cycles was achieved by exposing the tissue to a bleaching solution containing 4.5% hydrogen peroxide and 24 mM sodium hydroxide under LED illumination for approximately one hour.

#### Image processing and quality control

Image processing was carried out using the containerized NextFlow workflow MCMICRO (https://github.com/labsyspharm/mcmicro).^101^ Illumination non-uniformity was corrected using BaSiC flat-fielding.^102^ For each specimen, images from all staining cycles were computationally stitched and spatially aligned into a single multiplexed image using ASHLAR.^103^ The resulting multichannel composites were then prepared for single-cell analysis. Nuclear probability maps were generated with the UnMicst v1 deep-learning model,^104^ and cell boundaries were delineated with the S3segmenter marker-controlled watershed algorithm (https://github.com/HMS-IDAC/S3segmenter). Segmentation outputs were subsequently quantified with MCQuant (a module for single cell quantification in MCMICRO), which measured marker intensities for every cell across the entire slide (https://github.com/labsyspharm/quantification). To ensure high-quality data, staining patterns for each antibody were manually reviewed across all samples prior to downstream analysis. Additional image quality control was performed using the interactive tool CyLinter, which was used to tag and remove cells lacking detectable nuclei, cells lost across imaging cycles, and cells with aberrant marker expression profiles.^59^

#### 3D registration of CyCIF and H&E serial sections

To enable three-dimensional reconstruction of tissue architecture, serial CyCIF sections from Patient1_B4 were co-registered at full spatial resolution using a multistage registration strategy optimized for histological tissue deformation. Nuclear Hoechst staining, present in every CyCIF cycle, was used as the alignment channel to ensure robust correspondence of cellular and structural features across adjacent sections. Registration was performed pairwise between consecutive sections using Advanced Normalization Tools (**Figure S3A**).^54,55,105^ Each registration began with center-of-mass alignment, followed by rigid and affine transformations to correct for global translation, rotation, scaling, and shear. Nonlinear deformation was then modeled using the symmetric diffeomorphic normalization (SyN) algorithm to correct for local distortions. Mutual information was used as the similarity metric throughout all stages to accommodate intensity variations across sections and staining cycles. A multi-resolution optimization scheme was employed, with progressively finer image downsampling and smoothing levels, to improve convergence and robustness. Composite transforms were applied to resample each section into a common reference space, preserving full image resolution for downstream analyses. To generate a consistent 3D tissue volume, all registered CyCIF sections were stacked in their physical z-order using the known section thickness and inter-section spacing. The resulting 3D reconstruction enabled accurate spatial localization of cells, tissue landmarks, and higher-order structures such as granulomas and tertiary lymphoid structures across serial sections (**Figures 1J, S3D; Video S1**). More details and code can be found as listed in the software availability section.

H&E-stained serial sections from the same tissue block were independently co-registered using an elastix-based pipeline optimized for histopathology images.^56,106^ Similar to the CyCIF workflow, registration proceeded through rigid and affine alignment followed by nonlinear deformation, using mutual information as the similarity metric. This ensured accurate spatial correspondence across H&E sections while accommodating staining-specific contrast and tissue deformation (**Figure 1G**). Registered H&E volumes were used for complementary histomorphological analyses and cross-modality comparisons with CyCIF-derived reconstructions. Registered CyCIF (and H&E) image volumes were imported into Imaris (v10.2.0; Oxford Instruments, formerly Bitplane) for three-dimensional visualization, analysis, and surface generation.

#### Manual and automated annotation of three-dimensional tertiary lymphoid structures

Annotation of tertiary lymphoid structures (TLSs) was performed on CyCIF serial sections. CD20⁺ aggregates (with associated CD3E⁺ cells) were manually annotated in the first CyCIF slide (Section_2, z = 5 µm). QuPath (v0.3.2, v0.5.1, v0.6.0) was used for immunofluorescence-based automated annotation of TLS regions. A thresholder-based pixel classifier was applied to segment CD20⁺ TLSs, followed by dilation of annotations to capture peripheral TLS T cell zones (**Figures 1H, S1D**). Manual review and adjustment of the resulting regions of interest (ROIs) were performed in OMERO PathViewer™ 3.7.1 (Glencoe Software Inc.), with images hosted on the OMERO server (OMERO.server version 5.6.6-132-gs-ice36, accessed through OMERO.web 5.22.1; Glencoe Software Inc., University of Dundee & Open Microscopy Environment).

To segment 3D TLS objects from 2D annotations across serial sections, connected-component analysis was performed on binarized TLS ROI masks in MATLAB (R2022a) (**Figure 1I**). Discrete 3D TLS structures were identified using 26-connectivity to link voxels across adjacent sections. Labeled components were generated with *bwlabeln*, and morphological features – including volume, centroid coordinates, and shape descriptors – were quantified using *regionprops3*. Automated segmentations were subsequently reviewed in the MATLAB Volume Segmenter app (Image Processing Toolbox v11.5), where manual curation corrected instances of over-merging or over-fragmentation by splitting erroneously fused TLSs and merging components representing a single structure.

For downstream analyses, the curated 3D TLS segmentations were exported as a labeled 3D TIFF mask registered to the CyCIF image volume. Each cell in the single-cell CyCIF dataset was spatially queried against the 3D TLS mask, and cells located within TLS volumes were annotated with a TLS ROI label or ID derived from the MATLAB-generated mask. For visualization and additional image-based analyses, the 3D labeled mask was imported into Imaris to render the 3D TLSs as surface representations in the tissue image volume (**Figures 1J**, **S3D**, **2A-D; Video S1**).

In total, 906 3D TLS objects were segmented. The three-dimensional completeness of individual TLSs was evaluated using a custom ImarisXT Python script (https://github.com/labsyspharm/TB_3D_TLS_2026/). For each TLS surface object, an axis-aligned bounding box was computed from the centroid position and the Extent (BoundingBoxAA Length) statistics. These bounding boxes were then compared with the physical bounds of the registered image volume derived from the dataset dimensions and voxel spacing. TLS objects whose bounding boxes were fully contained within the X–Y–Z limits of the image volume were classified as complete, whereas objects whose bounding boxes intersected any image volume boundary were classified as incomplete or partially sampled. Using this criterion, 640 TLS objects were classified as complete and 266 as partial.

In the serial-section reconstruction dataset (Patient1_B4), specific markers were excluded because staining failed in some samples: Section_53 (CD4, CD11c, PDL1, PCNA), Section_98 (CD8a, TIGIT, PD1, αSMA), and Section_104 (PNAd, FOXP3, CD21). Corresponding cell assignments or phenotypes dependent on these markers were therefore excluded from downstream analyses. For single-cell image data analysis, more stringent quality control was applied. Only TLSs in which ≥75% of constituent cells passed previously described single-cell QC filters were retained, yielding 470 TLSs for downstream analyses. For optimal transport modeling and OT-based analyses, we further required TLSs to (i) span three or more serial sections and (ii) pass visual assessment of 3D structure and continuity, resulting in 366 TLSs.

Observed discontinuities in tissue morphology from one section to the next in the final image suggested that section order might have been slightly shuffled or a section lost in one or two locations in the stack. This type of problem arises during sectioning and reflects the challenge of working with archival tissue specimens; in the current paper we used the nominal section order because there was no objective means of correcting it.

Note that the total number of cells in each 3D TLS segmentation mask (generated by connected component analysis as illustrated in **Figure 1I**) is equal to the number of successfully segmented cells on 2D sections lying within that mask. Because CyCIF images are separated by two 5-µm tissue sections that did not contribute to these values, the actual number of cells in 3D objects is likely higher than the recorded values by ∼2 fold (based on an average 10 µm cell diameter); no correction was made for this undercount. See Yapp et al.^48^ for further discussion of this point.

#### Quantitative single-cell image analysis

Using the open-source interactive gating platform Minerva Analysis Gater (https://github.com/labsyspharm/minerva_analysis), single-cell marker intensities were thresholded to generate binary gates. Single-cell image quantification and data preprocessing were performed using the SCIMAP Python package.^107^ Marker-specific gating thresholds were used to normalize single-cell intensity measurements to a 0–1 scale, and values above 0.5 were classified as marker-positive. Cell-type annotation was then carried out using gating strategies adapted from validated flow-cytometry panels encompassing a broad range of immune and stromal populations (**Figure 1E**). The primary phenotyping panel included CD20, CD3E, CD4, CD8a, FOXP3, CD11c, CD206, CD86, αSMA, pan-CK, CD31, MCT, and MPO (**Figure S2C**).

Additional sub-phenotyping schemes were applied as follows: Ki-67 was used to identify proliferating cells; CD21 and CD23 were used to classify follicular structures into primary (CD21⁺) and secondary (CD21⁺CD23⁺) follicles, as well as proliferating germinal centers (Ki-67⁺CD21⁺CD23⁺). TIGIT expression in T cells proximal to follicles was used to define follicular T cell subsets, including T follicular helper cells (Tfh; TIGIT⁺ CD4 T cells), T follicular regulatory cells (Tfr; TIGIT⁺ Tregs), and follicular cytotoxic T cells (Tfc; TIGIT⁺ CD8 T cells). PNAd⁺ subsets of CD31⁺ endothelial cells were profiled as high endothelial venules (HEVs) components. Spatial LDA was performed using the SCIMAP function (*scimap.tl.spatial_lda*), with the kNN method and k = 10.

#### kNN-based definition of functional zones within 3D tertiary lymphoid structures

TLS zones were defined based on the spatial organization of canonical B cell and FDC markers using K-Nearest Neighbors (KNN) algorithm.^108^ Seed zones were first generated from CD20, CD21, or CD23 single cells, yielding CD20⁺ B cell populations, CD21⁺ follicular dendritic cell–rich regions, and CD21⁺CD23⁺ double-positive regions within each TLS. These seeds correspond to expected architectural subzones: CD21⁺CD23⁺ zones being germinal centers, CD21⁺ regions representing follicles (including primary follicles and secondary follicles, if containing a GC), and CD20⁺ B cell zones. Zone masks were combined into mutually exclusive labels using the following priority: CD21⁺CD23⁺ > CD21⁺ > CD20⁺. Voxels not assigned to any of these zones were categorized as “Other” (i.e., the rest of TLS).

These initial zones were used for TLS zone volume proportion and size analyses (**Figures 3F-G**). Because single-cell positivity alone yielded sparse and discontinuous label maps, seed sets were converted into continuous regional zones. Binary masks derived from marker-positive cells were radially expanded in the XY plane only (i.e., within individual sections, preserving section thickness) using a 50-pixel (≈16.25 µm) 2D Euclidean expansion kernel. This procedure captures cells immediately adjacent to highly positive cores and better approximates anatomically continuous follicular regions without artificially merging structures across sections. The resulting labeled TLS zones were used for downstream analyses of zone-specific cellular composition and spatial organization.

#### Optimal transport-based modeling of TLS maturation trajectories

To model TLS maturation as a continuous process rather than a discrete set of classes, we applied an optimal transport (OT) framework. We assumed that TLSs with similar spatial arrangements of cell phenotypes occupy nearby positions along a shared developmental continuum. Accordingly, pairwise unbalanced fused Gromov-Wasserstein (UFGW) distances were used to quantify the spatial-phenotypic remodeling required to transform one TLS into another, such that TLS pairs separated by a low UFGW distance were interpreted as developmentally closer, whereas those separated by high UFGW distance were interpreted as farther apart.

Directed UFGW distances were computed using an entropy-regularized, low-rank solver in OTT-JAX (**Figure 4A**). Since TLS size is a strong covariate of structural organization (**Figure 3B**), anchor TLSs were selected to span the full-size range of the ensemble. Distance profiles from each TLS to these anchors were then embedded via a Deep Set encoder and projected into diffusion-map space, with directed progression inferred by flow matching. Corresponding flow time or pseudotime were derived accordingly from each trajectory (**Figures S6A-C**). Full mathematical details and parameter settings are provided in the Supplementary Methods.

#### Spatial Transcriptomics by GeoMx digital spatial profiling

Freshly cut 5-µm serial sections from 3 patient samples (Patient1_B5_Section_2, Patient2_Section_12 and Patient3_Section_12) were processed for GeoMx spatial transcriptomics, as described by the manufacturer (Bruker, formerly NanoString Technologies).^109^ Guided by immunofluorescence staining (DNA-FITC/525 nm, pan-cytokeratin-Cy3/568 nm, CD45-Texas Red/615 nm, CD20-Cy5/666 nm), a total of 161 histopathological regions of interest (ROIs) were selected for Human Whole Transcriptome Atlas (WTA) profiling, including 133 TLS ROIs, 7 lymph node ROIs (LN, all from Patient2), and 21 uninvolved lung ROIs (UL, negative control for lung alveoli). The library was then sequenced with Illumina NovaSeq 6000. Quality control and quartile-3 (Q3) normalization of the dataset were then performed as previously detailed.^110^ (https://www.bioconductor.org/packages/release/workflows/vignettes/GeoMxWorkflows/inst/doc/GeomxTools_RNA-NGS_Analysis.html). 16 ROIs from Patient3 (10 TLS, 6 UL ROIs) were excluded from subsequent analysis due to an unusually low number of sequencing reads, consistent with poor tissue quality. Analyses were performed in R (v4.5.1) using RStudio (v2024.12.0), including dimensionality reduction and clustering analyses using UMAP (v0.2.10.0). Differential expression was assessed using DESeq2 (v 1.48.1). After GeoMx spatial profiling, same-slide CyCIF was performed on a subset of tissue sections to phenotypically deconvolute the GeoMx regions of interest.

In parallel to OT analysis on 3D image data, pseudotime analysis of GeoMx transcriptomic profiles derived from 2D TLS regions was performed to identify progressive transcriptional programs associated with TLS development FDC differentiation (**Figures S1E-G**). GeoMx ROI expression profiles were log1p-normalized and reduced by PCA (10 components using the top 200 highly variable genes). A diffusion map embedding was computed using a self-tuning Gaussian kernel with adaptive bandwidth and symmetric Markov normalization. Pseudotime was defined along DC1, rooted at the Early TLS ROI with the minimum DC1 value, and rescaled to [0, 1]. A principal curve (cubic smoothing spline) was fitted to visualize the trajectory. The top 200 genes were selected by coefficient of variation (CV = σ/μ). Z-scored expression (clipped to ±2) was plotted with ROIs ordered by pseudotime and genes ordered by peak pseudotime (**Figure S1F**). Overlay lines represent the mean z-score of genes positively and negatively associated with pseudotime (Spearman correlation with BH-FDR adjustment). Spearman correlation between gene expression and pseudotime was computed across all ROIs and adjusted using BH-FDR.

#### TIGIT⁺ CD4 T cell and Treg abundance analysis

TLSs were stratified based on the joint distribution of TIGIT⁺ CD4 T helper (Th) cells and TIGIT⁺ regulatory T cells (Tregs). For each TLS, a relative enrichment metric was defined as *r* = (TIGIT⁺ Treg) / (TIGIT⁺ Th cell + 1), where a pseudocount was included to stabilize low-abundance estimates. TLSs were partitioned into four quadrants (Q1–Q4) using thresholds derived from cohort-wide distributions. The median abundance of TIGIT⁺ Th cells defined the vertical boundary (x_thr), and the mean ratio of TIGIT⁺ Treg-to-TIGIT⁺ Th cell defined a nonlinear decision boundary in log₁₀ space: log₁₀(1 + TIGIT⁺ Treg) = log₁₀[r_thr × (TIGIT⁺ Th cell + 1)]. These boundaries defined four quadrants: Q1, low TIGIT⁺ Th cell abundance with above-average Treg enrichment; Q2, low TIGIT⁺ Th cell abundance with above-average Th enrichment; Q3, high TIGIT⁺ Th cell abundance with above-average Th enrichment; Q4, high TIGIT⁺ Th cell abundance with above-average Treg enrichment. Thresholds and 95% confidence intervals were estimated by bootstrap resampling (n = 500). The relationship between TIGIT⁺ Th cells and TIGIT⁺ Tregs was quantified using ordinary least squares regression in log₁₀-transformed space: log₁₀(1 + TIGIT⁺ Treg) = a + b × log₁₀(1 + TIGIT⁺ Th cell), with goodness-of-fit assessed by R² and Spearman correlation. TLS cluster identities were overlaid onto the quadrant embedding for downstream analyses, and kernel density estimation was used to visualize quadrant distributions.

#### Unsupervised feature discovery and automated pixel-level segmentation of lymphoid aggregates from H&E images

To enable automated and unbiased detection of lymphoid aggregates in H&E serial sections, we applied a previously-described unsupervised histomorphology feature discovery framework.^111^

#### 2D TLS age score (tₘ) estimation

For each 2D TLS section, a Voronoi tessellation was computed from 2D cell coordinates. Cells sharing a tile edge were connected to form an adjacency graph. Breadth-first search (BFS) over this graph up to five hops yielded exclusive rings. Per-hop features included ring cell counts, mean intercellular distances, and B-lineage/non-B cell fractions. At hop 1, the mean distance to each cell phenotype was additionally recorded. At hops 3 and 5, the distance to the neighborhood centroid captured the spatial eccentricity of each cell within its broader microenvironment. Spatial features were aggregated per slice (mean, standard deviation, IQR), and a HistGradientBoosting Model (HistGBM) regressor was trained using 5-fold cross-validation on an 80/20 TLS split to predict the 3D tₘ assigned to each slice.

#### Prediction of 3D tₘ from consecutive 2D slices

Summary statistics of the predicted 2D TLS age score across *n* sampled slices (mean, standard deviation, quartiles, slice count) were used to train a HistGBM regressor to predict the corresponding 3D tₘ. To simulate limited data, n = 1–15 consecutive slices were bootstrap-sampled (500 draws per TLS) and evaluated against the full-stack reference by Spearman ρ and R².

## ACKNOWLEDGMENTS

This work was primarily funded by a grant from the Gates Foundation INV-027106 (PKS, BA). Additional support was provided by the Harvard Ludwig Center (for NSCLC imaging) and NIAID R21 AI155003-01 (PKS, BA, AJM), the Global Research Resource for Human Tuberculosis (https://sites.uab.edu/tbresearchresource/; NIH R24AI186591); the Wellcome Trust (Africa Health Research Institute strategic core award 227167/A/23/Z, and Discovery Award 314407/Z/24/Z). We thank members of the LSP spatial profiling team, particularly B.K. Kobs, M. Lopez Leon and Y. Chen for expert assistance throughout the project.

## AUTHOR CONTRIBUTIONS

Study conceptualization and design: E.C.O., S.R.T., Z.M., and P.K.S. Imaging data collection and analysis: E.C.O., S.R.T., Z.M., Z.Z., C.Y., Y.D.L., and J.M. 3D image reconstruction and computational modeling: S.R.T. Tissue specimen collection, image review, and pathological annotation: A.G., T.N., A.J.M., S.S., A.R.S., and I.H.S. Supervision and funding: A.J.M., S.S., B.B.A., A.J.C.S., and P.K.S.

---

**Figure S1.**
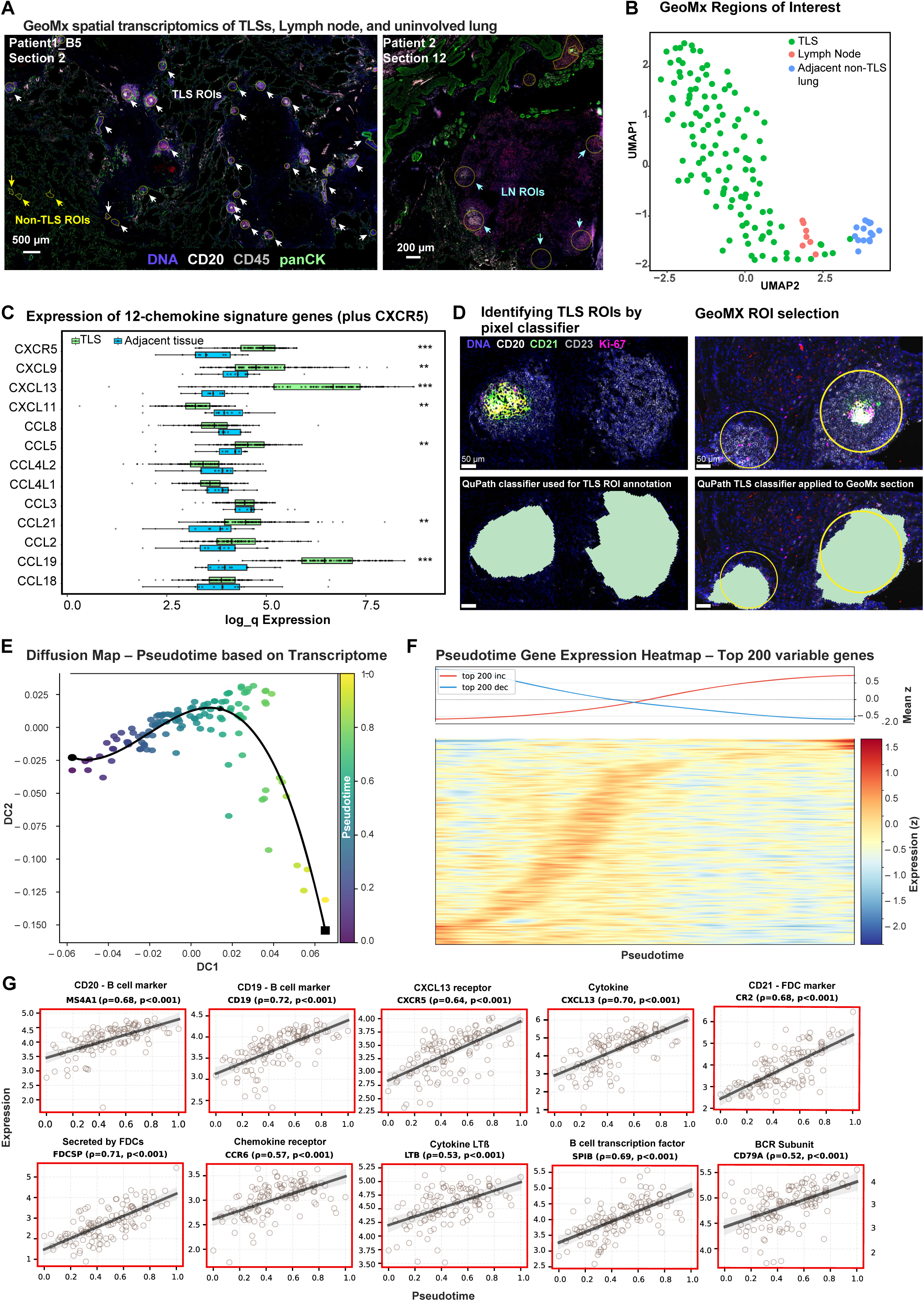
Spatial transcriptomic validation of TLS identity in TB-infected human lung tissue, related to Figure 1. **(A)** GeoMx spatial transcriptomics of tertiary lymphoid structures (TLSs), lymph node, and uninvolved lung alveoli from TB-infected human lung samples. Representative regions of interest (ROIs) are shown from Patient1_B5 (section 2) and Patient2 (section 12), overlaid on immunofluorescence images (DNA, CD20, CD45, pan-cytokeratin). **(B)** UMAP embedding of GeoMx ROIs colored by histological region, demonstrating transcriptional separation between lymphoid structures (TLSs, lymph node) and uninvolved lung regions. **(C)** Expression of members of the 12-chemokine (12-CK) TLS gene signature, together with CXCR5, in TLSs compared to uninvolved lung (UL) regions. Boxplots show log₂-transformed Q3-normalized expression values for individual regions of interest, with TLS regions (green) and UL regions (light blue). Boxes represent the interquartile range (25th–75th percentiles), the center line indicates the median, and points represent individual measurements. CXCR5 is shown for comparison but is not part of the canonical 12-chemokine TLS signature. **(D)** Cross-modal validation of TLS identification. Top left: Representative TLS from Patient1_B4 (used for 3D reconstruction) highlighting B-cell and follicular organization. Top right: Multiplexed CyCIF image from Patient1_B5 (section 2; post-GeoMx CyCIF) showing a TLS ROI with the indicated markers (DNA, CD20, CD21, CD23, Ki-67). Yellow outlines indicate annotated GeoMx TLS boundaries. Bottom panels: Application of a QuPath TLS pixel classifier to identify TLS regions in both the 3D reconstruction and GeoMx sections, demonstrating concordant detection across modalities and validating the annotated TLS ROIs in serial section reconstruction experiment as *bona fide* TLSs. **(E)** Diffusion map of 2D TLS transcriptional states using GeoMx ROIs, colored by transcriptomic pseudotime. Each point represents one ROI plotted in diffusion space (DC1, DC2). A principal curve (solid black line) indicates the inferred maturation trajectory. **(F)** Heatmap of the top 200 most variable genes (ranked by coefficient of variation) across 2D TLSs. Z-scored gene expression (± 2) is plotted along pseudotime. Overlay lines indicate average z-score trends of the top 200 most variable increasing (red) and decreasing (blue) gene sets along the pseudotime. **(G)** Signatures of TLS pseudotime based on transcriptome. Log₂-transformed Q3-normalized expression of selected genes along pseudotime. Each panel reports Spearman ρ and p-value.

**Figure S2.**
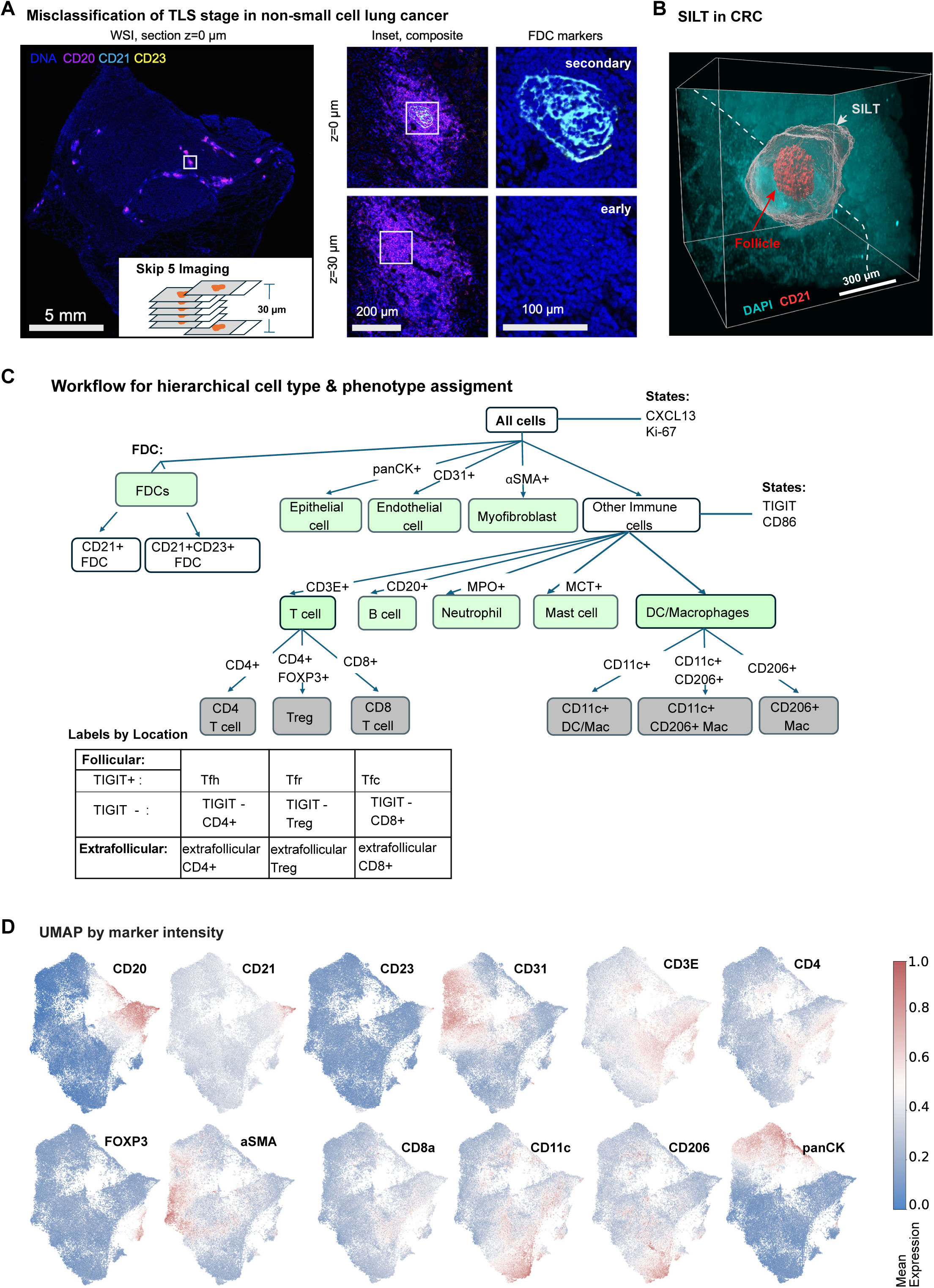
Examples highlighting limitations of 2D TLS classification in other tissues and the cell phenotyping strategy used for TB TLS profiling, related to Figure 1. **(A)** TLS misclassification observed in non-small cell lung cancer tissue sample. **(B)** 3D image of representative solitary intestinal lymphoid tissue (SILT) in colorectal cancer tissue. The CD21⁺ follicular dendritic cell (FDC) network occupies only ∼8.4% of the total SILT volume, highlighting the potential for misclassification in 2D tissue sections if the follicular compartment is not captured. **(C)** Phenotyping workflow for assigning cell types. **(D)** Uniform manifold approximation and projection (UMAP) plot of single-cell data from CyCIF imaging (Patient1_B4_Section 2), colored by marker intensity. UMAP was generated from randomly sampled 50,000 single cells. See Figure 1F.

**Figure S3.**
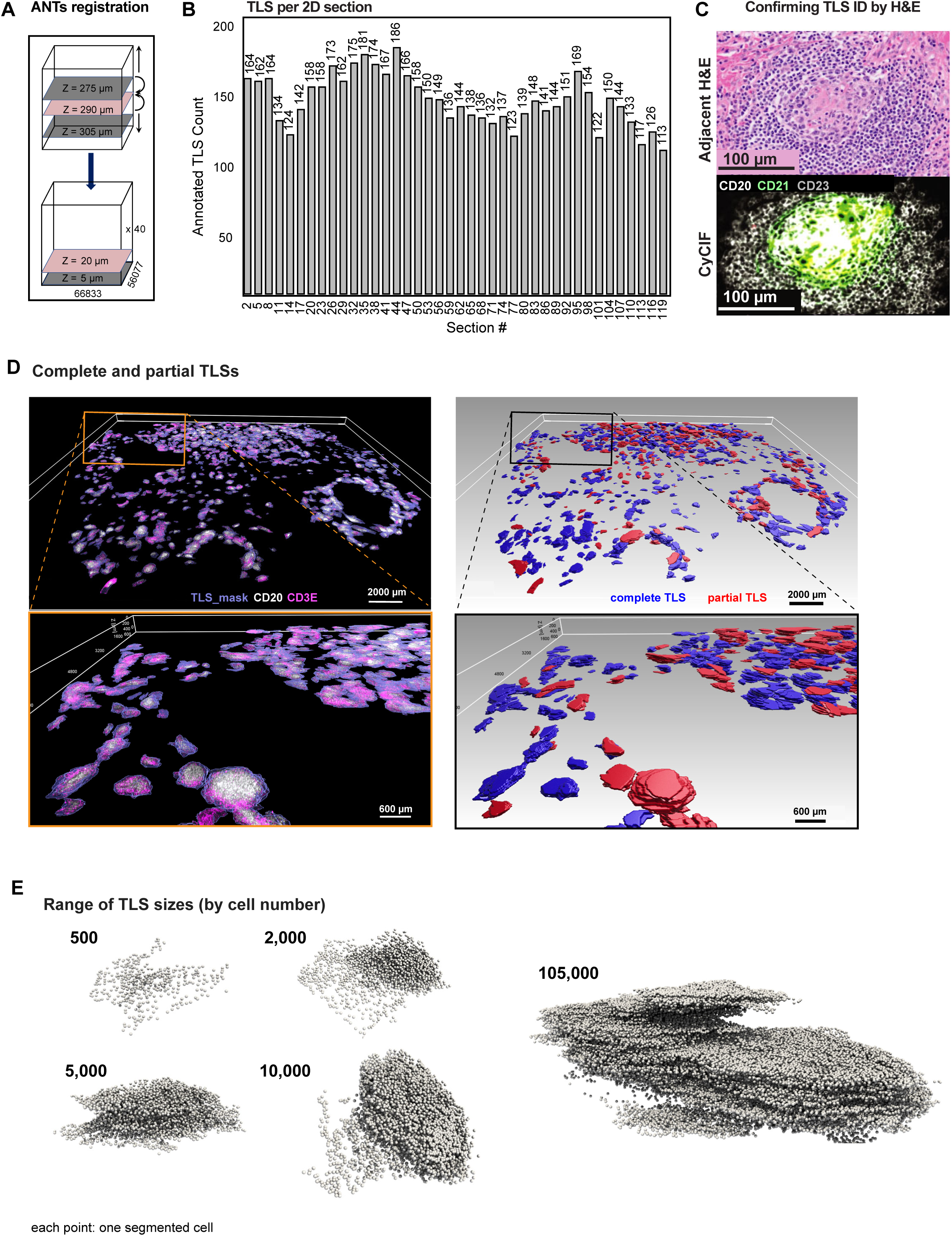
Three-dimensional reconstruction reveals heterogeneous TLS architectures in pulmonary TB, related to Figure 1. **(A)** Representation of ANTs-based registration pipeline for CyCIF serial section alignment to generate a spatially coherent 3D tissue volume. **(B)** Number of annotated TLS regions per serial section across Patient1_B5, illustrating section-to-section variability in TLS detection, motivating volumetric reconstruction. **(C)** Representative TLS ROIs showing concordant H&E morphology and CyCIF marker expression. H&E-to-CyCIF registration enabled comparison between annotation of TLSs by both modalities. Scale bars, 100 µm. **(D)** Assessment of volumetric completeness of TLSs in TB-infected lung. Right panels: Whole-slide and higher-magnification CyCIF images showing TLS masks (blue) with CD20 (white) and CD3E (magenta) staining. Left panels: Corresponding surface-rendered 3D reconstructions of TLS objects generated from serial sections. TLSs fully contained within the reconstructed image volume were classified as complete (blue), whereas TLSs truncated by volume boundaries were classified as partial or incomplete (red). **(E)** Representative 3D renderings of TLSs spanning the size range in dataset, from small aggregates to very large extended structures. Cells are rendered in ParaView.

**Figure S4.**
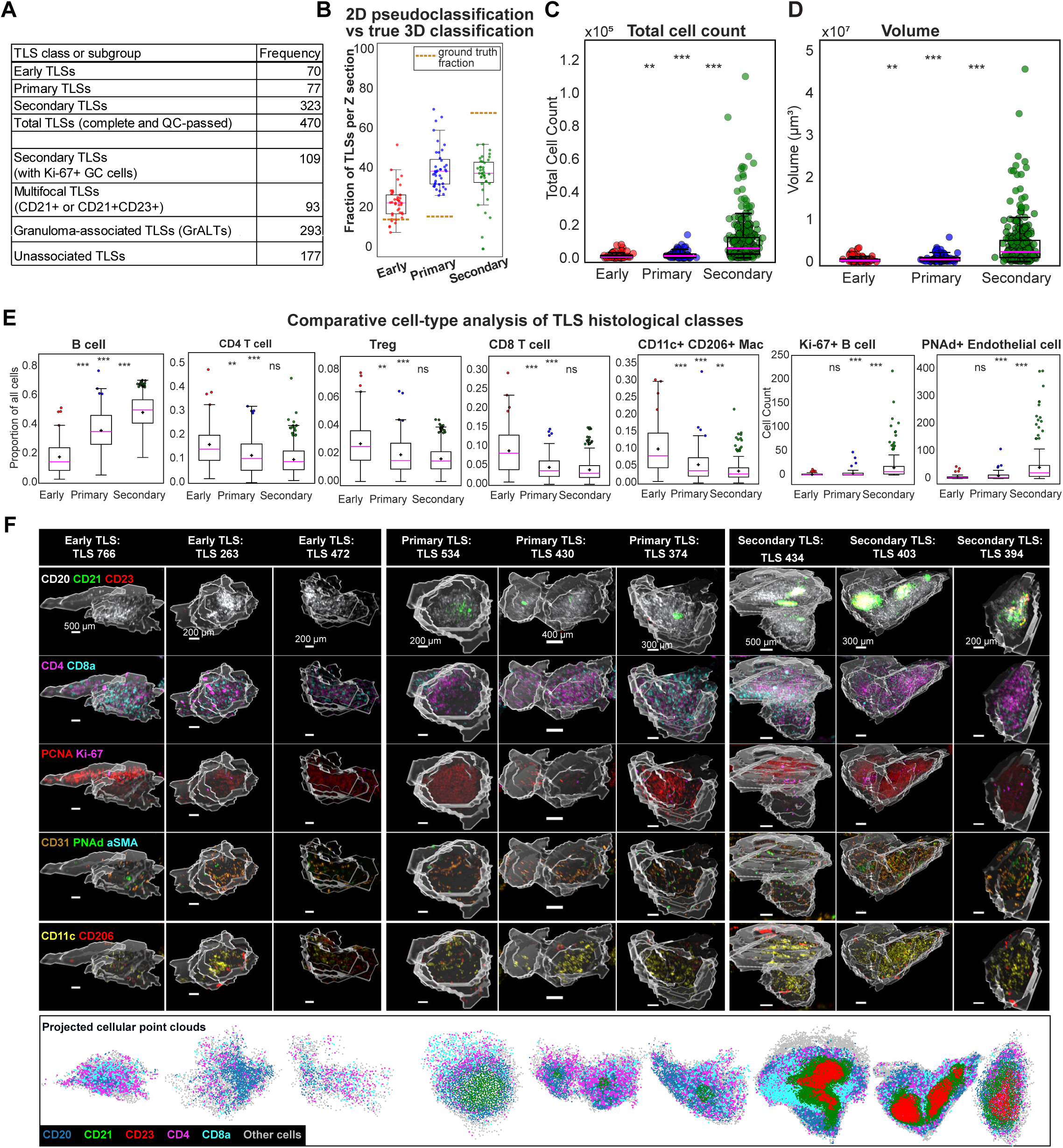
Three-dimensional characteristics of canonical TLS maturation stages, related to Figures 2 and 3. **(A)** Summary of 3D TLS classes identified in the reconstructed Patient1_B4 tissue image. 3D TLSs were categorized as Early, Primary, or Secondary based on CD20⁺ B cells and CD21⁺ and CD23⁺ follicular dendritic cells (FDCs). Counts are shown for complete TLSs that passed QC, including subsets exhibiting Ki-67⁺ germinal center (GC) cells, multifocal organization, and spatial association with granulomas. **(B)** Per-section pseudoclassification versus true 3D TLS classification. Burnt orange dashed lines indicate true class proportions across 470 TLSs (see **Figure S4A**). **(C-D)** Quantification of TLS size and volume across histological classes. Total cell count (B) and TLS volume (C) increase progressively from Early to Primary to Secondary TLSs, indicating coordinated structural expansion with maturation. Each point represents a single TLS. **(E)** Comparative cell-type composition across TLS classes. Boxplots show relative proportions or absolute counts of major immune and non-immune populations. Secondary TLSs exhibit increased levels of total and Ki-67⁺ B cells and PNAd⁺ HEVs relative to Early and Primary TLSs. Proportionally, other cell types like CD4⁺ and CD8⁺ T cells, Tregs, CD11c⁺ CD206⁺ macrophages decline across maturation stages, reflecting relative contraction of these compartments as the B cell and follicular compartments expand. Center line, median; box, interquartile range; whiskers, 1.5× IQR. Top 5% of values are shown as jittered points; black crosses indicate means. Statistical significance was assessed by pairwise Mann-Whitney U tests (*P* < 0.05; \**P* < 0.01; \*\**P* < 0.001). **(F)** Representative three-dimensional renderings of TLSs across maturation classes (early, primary, and secondary). Rows show marker groups highlighting B cell and follicular organization (CD20, CD21, CD23), T cell subsets (CD4, CD8a), proliferative activity (PCNA, Ki-67), vascular and stromal features (CD31, PNAd, αSMA), and myeloid populations (CD11c, CD206). These examples illustrate progressive follicular organization, immune compartmentalization, and proliferative activity with TLS maturation, while also highlighting intra-class heterogeneity. **Bottom:** Corresponding 2D projections of cellular point clouds for each TLS, generated by centering and principal axis alignment, with cells colored by phenotype to visualize spatial organization of B cells, T cells, and follicular domains.

**Figure S5.**
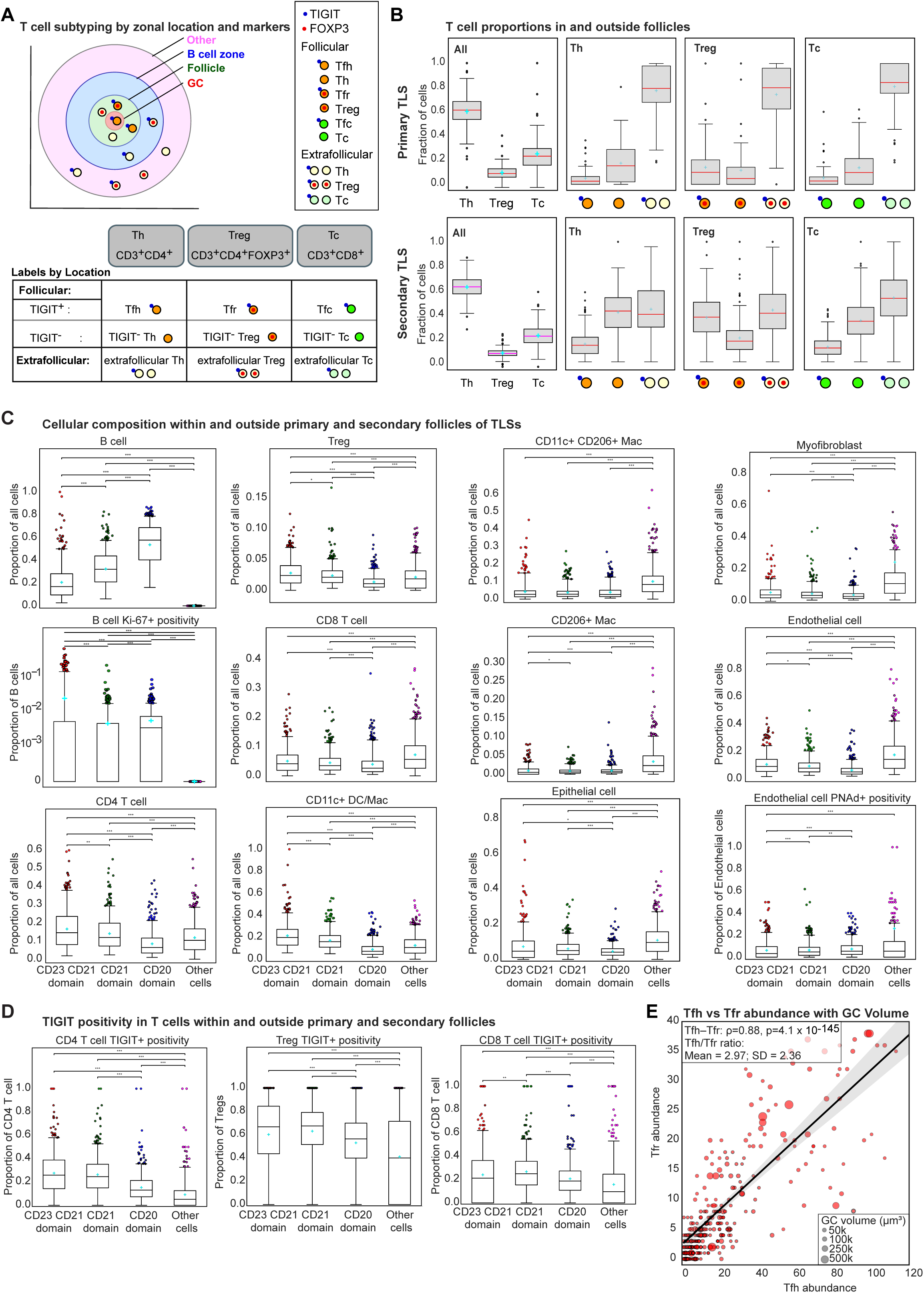
Spatial organization of immune cells across follicular and extrafollicular TLS compartments, related to Figure 3. **(A)** T cell subtyping by zonal location and markers. TLS schematic showing concentric shells based on mean volume-equivalent radii of TLS zones. The spatial positioning of follicular and extrafollicular T cells are also indicated. Also shown is the phenotyping scheme used to determine follicular and extrafollicular T cell states. We defined T follicular helper cells (Tfh) cells as CD3^+^ CD4^+^ TIGIT^+^ FOXP3^-^ in the follicle or GC; T follicular regulatory cells (Tfr) cells were similar but also FOXP3^+^; and TIGIT^+^ CD3^+^ CD8^+^ T cells in GC or follicle were follicular cytotoxic T (Tfc) cells. TIGIT was used for quantification instead of PD1 because it had better signal to noise ratio. There were T cells in the follicle or GC that did not express TIGIT. T cells outside the follicle whether expressing TIGIT or not were defined as extrafollicular T helper (Th), Tregs, or cytotoxic T cells (Tc). **(B)** Fractional composition of total and spatially-resolved T cell subsets within primary and secondary TLSs. First panels: fraction of CD4⁺ T cells, Tregs, and CD8⁺ T cells relative to their sum per TLS. 2nd-4th panels: fractional composition of spatially-resolved subsets of CD4⁺ T cells, Tregs, and CD8⁺ T cells. Boxplots show the distribution of per-TLS fractions, with cyan crosses representing means, center lines indicating medians, boxes representing interquartile ranges, whiskers denoting 1.5× IQR, and points representing individual TLSs. **(C)** Quantification of cellular composition within and outside primary and secondary follicles. Boxplots show proportions of major immune, stromal, and structural cell types across CD20⁺, CD21⁺, CD23⁺CD21⁺, and “Other” regions, revealing strong zone-specific enrichment patterns. Center line, median; box limits, upper and lower quartiles, whiskers extend at most to 1.5x IQR; cyan crosses indicate means. Statistical comparisons were performed using pairwise Mann-Whitney U tests (P < 0.05; *P < 0.01; **P < 0.001). **(D)** TIGIT positivity across T cell subsets stratified by spatial zones. Boxplots show the fraction of TIGIT⁺ cells among CD4⁺ T cells, Tregs, and CD8⁺ T cells within follicular (CD21⁺ and CD23⁺CD21⁺), CD20⁺, and “Other” regions, demonstrating preferential enrichment of TIGIT⁺ T cells within follicular compartments. **(E)** Tfh abundance positively correlates with Tfr abundance across CD21⁺CD23⁺ GC domains. Here, abundance refers to cell counts. Spearman correlations: Tfh–Tfr, ρ = 0.88 (p = 4.1 × 10⁻¹⁴⁵); Tfh:Tfr ratio–GC volume, ρ = 0.05 (p = 0.36). Mean Tfh:Tfr ratio = 2.97; SD = 2.36. Of the 435 GC domains, 338 (77.7%) contained both Tfh and Tfr cells and only these were included in ratio analyses. Each point represents a GC domain and bubble size reflects 3D GC volume. Black line indicates linear regression with 95% confidence interval shading.

**Figure S6.**
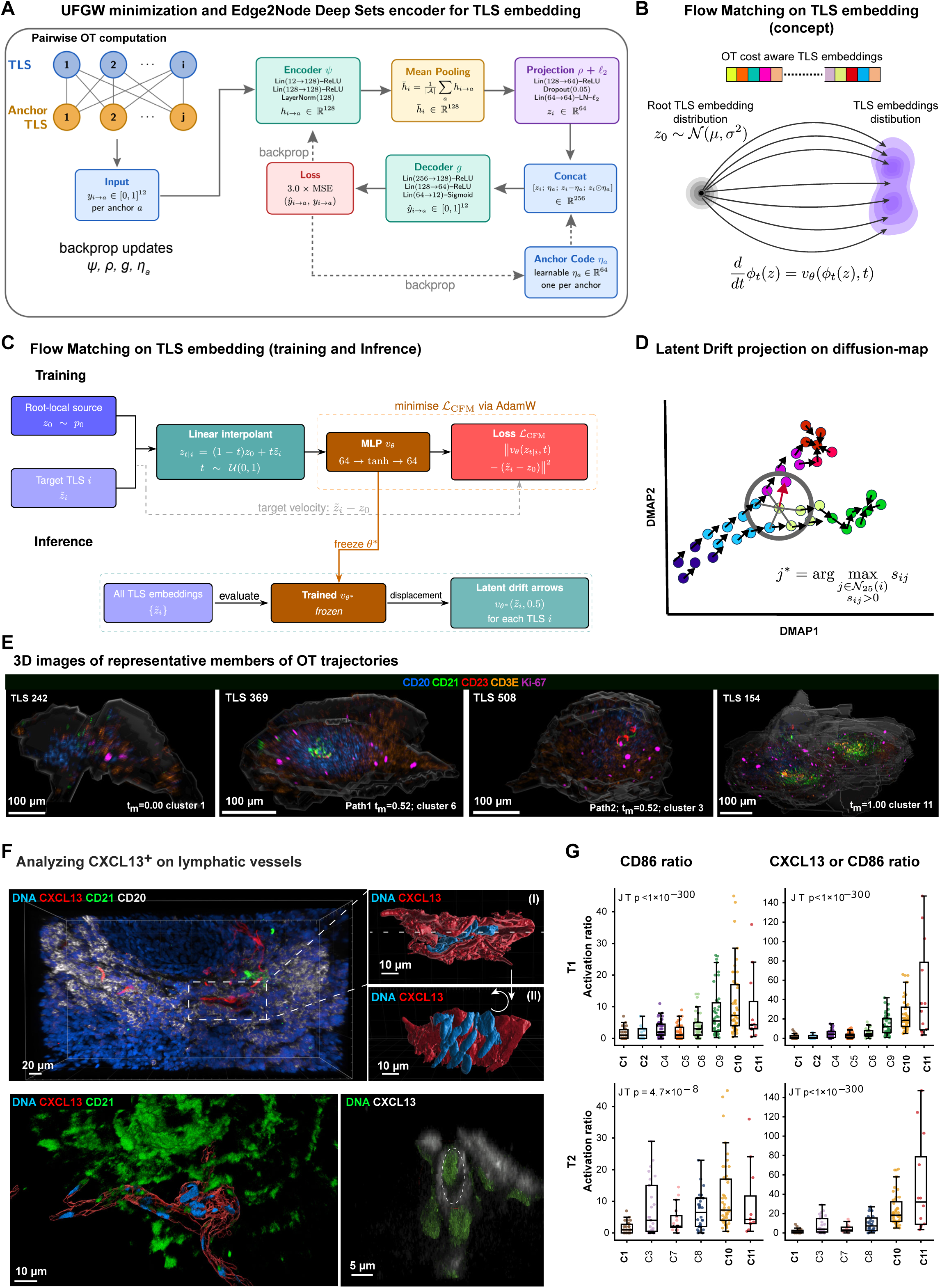
Maturation trajectory inference and follicular activation in TLSs, related to Figures 4 and 6. **(A)** UFGW minimization and Edge2Node Deep Sets encoder for TLS embedding. Pairwise transport-cost vectors between each TLS and a set of anchor TLSs were computed using Unbalanced Fused Gromov-Wasserstein (UFGW) minimization. UFGW jointly aligns the internal 3D spatial geometry of each TLS and penalizes phenotype mismatches across super-cells. It accommodates differences in total cell count via KL-penalized mass creation and removal terms. Each normalized edge vector *y*_*i*_ → *a* ∈ [0,1]^12^ encodes displacement, birth-cost, and death-cost across phenotypes. A shared encoder *ψ* maps each edge vector to a 128-dimensional hidden representation ℎ_*i*_ → *a*. These representations are aggregated by mean pooling over all anchors to yield a permutation-invariant node summary ℎ_*i*_ ∈ ℝ^128^. A projection head ρ followed by ℓ₂ normalization produces the final TLS embedding *z*_*i*_ ∈ ℝ^64^. During training, a decoder g reconstructs each edge vector *ŷ*_*i*_ → *a* from *z*_*i*_ concatenated with a learnable anchor code *η*_*a*_, minimizing a reconstruction loss via backpropagation through *φ*, *ρ*, *g*, and *η*_*a*_. After training, *z*_*i*_ serves as the compact, permutation-invariant transport-cost embedding for each TLS, used in all downstream diffusion-map and trajectory analyses. For details, see supplementary method. **(B)** Flow Matching on TLS embeddings (concept). OT-cost-aware TLS embeddings define a target distribution over the latent space (purple cloud, right). A root TLS source distribution *z*_0_∼*N*(*μ*, *σ*_0_^2^) is centered on the root TLS embedding (grey cloud, left). Flow matching learns a time-dependent vector field *v_θ_* satisfying the probability flow ODE 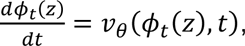 which continuously transports probability mass from the root distribution toward the target TLS distribution, representing a directed developmental drift across the latent space. **(C)** Flow Matching training and inference pipeline. A flow matching model is trained to learn a latent vector field *v*_*θ*_ that maps each TLS embedding toward its most probable next state. To this end, for every *TLS*_*i*_ a source point *z*_0_∼*N*(*μ*, *σ*_0_^2^) is sampled and a linear interpolant *z*_*t*|*i*_ = (1 − *t*)*z*_0_ + *tz̃*_*i*_ is constructed at uniformly sampled *z*_0_∼*U*(0,1). A time-conditioned MLP is trained to predict the conditional velocity *z̃*_*i*_ − *z*_0_ along the probability path, minimizing ℒ_*CFM*_ via AdamW for 10,000 epochs. Once trained, the model is used to infer displacement vector indicating the next most probable state in the latent space (Latent drift arrows). For details, see Supplementary Methods **(D)** Latent drift projection onto the diffusion map manifold and identifying the next most probable TLS state. The latent drift vector is projected onto the diffusion-map manifold using the Nyström extension approach (see Supplementary Information). The resulting displacement is then used to calculate an alignment score *S*_*ij*_ = *u*_*i*_ · *d*_*ij*_ for each of the *k* = 25 nearest neighbors in DMAP space. The neighbor with the highest positive alignment score, 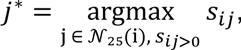 identifies the most probable next developmental state and defines the direction arrow visualized on the diffusion map manifold. **(E)** Representative 3D renderings of TLSs sampled along T1 and T2 at comparable flow times: TLS 242 (tₘ = 0.00), TLS 369 (tₘ = 0.52 on T1), TLS 508 (tₘ = 0.52 on T2), and TLS 154 (tₘ = 1.00). **(F)** CXCL13⁺ vessel-like structures show nuclear morphologies consistent with lymphatic endothelial vessels. Top left panel: Volumetric rendering of a TLS imaged with 3D CyCIF (DNA (blue), CXCL13 (red), CD21 (green), CD20 (white)). Scale bar is 20 µm. Top right panel (I): Zoomed-in view of a CXCL13⁺ vessel from top left panel (dashed box) and represented by surface rendering. Channels shown are DNA (blue) and CXCL13 (red). Scale bar is 10 µm. Top right panel (II): Cutaway section along dashed line in middle top panel showing multiple elongated nuclei (blue) within CXCL13⁺ vessel. Scale bar is 10 µm. Bottom left panel: Volumetric rendering of a different TLS (DNA (blue), CXCL13 (red), CD21 (green). CXCL13⁺ vessel is represented as translucent surface rendering (red). Nuclei (blue) shown are in contact with inner wall of vessel. Scale bar is 10 µm. Bottom right panel: Cross section of a 3D CyCIF dataset showing Hoechst-stained nuclei (green) within CXCL13 vessel (grey). Dashed outline highlights a vascular lumen containing nuclei. Scale bar is 5 microns. **(G)** Activated FDC ratios relative to inactive FDCs increase across clusters along both paths. Activation ratios are shown for CD86⁺ only (left) and CXCL13⁺ or CD86⁺ FDCs (right). FDCs are defined as (CD21⁺ or CD23⁺) CD20⁻ CD3E⁻ cells. Left: ratio of CD86⁺ FDCs to (1 + CD86⁻ FDCs). Right: ratio of (CXCL13⁺ or CD86⁺) FDCs to (1 + CXCL13⁻ CD86⁻ FDCs). p-values are from the Jonckheere-Terpstra trend test.

**Figure S7.**
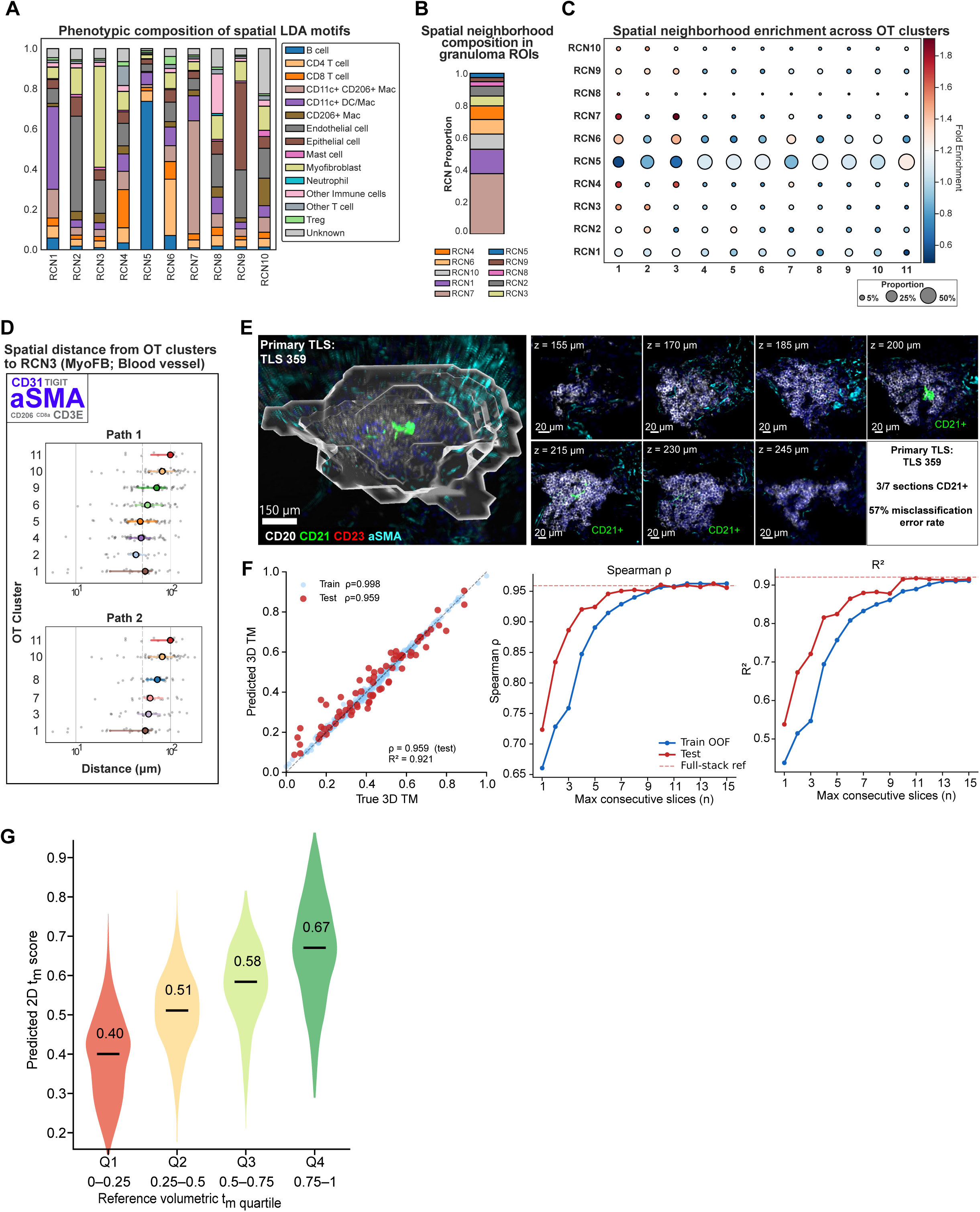
Spatial motif architecture and prediction of 3D TLS maturation state from 2D tissue sections, related to Figure 7. **(A)** Phenotypic composition of spatial motifs identified by latent Dirichlet allocation (LDA), representing recurrent cellular neighborhoods. **(B)** Motif composition within granuloma regions-of-interest (ROIs). **(C)** Motif enrichment across TLS OT clusters, with bubble size indicating motif proportion and color representing relative enrichment. **(D)** Distance from TLS OT clusters to RCN3 which corresponded to a CD31⁺ αSMA⁺ neighborhood. **(E)** Example of sectioning-based misclassification in a primary TLS. Individual serial sections variably display CD21 positivity, leading to apparent class switching in 2D. **(F)** Prediction of 3D TLS maturity (tₘ; mass-normalized time) from consecutive 2D sections using a gradient boosting model. Left: predicted versus true tₘ (train ρ = 0.998; test ρ = 0.959). Center/right: performance (Spearman ρ and R²) improves with increasing slice number, approaching full-stack accuracy at ∼11–13 sections. **(G)** Prediction of 3D TLS maturity from single 2D TLS sections. Predicted 2D tₘ scores stratified by reference 3D tₘ quartiles (Q1-Q4), showing a monotonic increase in predicted maturity. Prediction accuracy was highest for TLSs with intermediate reference 3D tₘ values (0.25–0.75), whereas highly mature TLSs showed greater variability, consistent with increased structural heterogeneity. See also Figure 7H.

